# A Multi-modal LLM-Knowledge Fusion Framework for Predicting Single-cell Genetic Perturbation Effects

**DOI:** 10.64898/2026.04.24.720560

**Authors:** Mingkun Lu, Nanxin You, Hongning Zhang, Lingyan Zheng, Bo Li, Wanghao Jiang, Yintao Zhang, Huaicheng Sun, Ying Zhou

**Affiliations:** College of Pharmaceutical Sciences, State Key Laboratory of Advanced Drug Delivery and Release Systems, Zhejiang University, Hangzhou, 310058, China; Department of Pharmacy, Second Affiliated Hospital, Zhejiang University School of Medicine, Hangzhou, 310009, China; Shenzhen Loop Area Institute, Shenzhen, 518038, China; Department of General Surgery, Sir Run Run Shaw Hospital, Zhejiang University School of Medicine, Hangzhou, 310016, China; Wisdom Lake Academy of Pharmacy, Xi’an Jiaotong-Liverpool University, Suzhou, 215123, China

**Author notes:** To whom correspondence should be addressed: Dr. Zhou. These authors contributed equally to this work as first authors (Mingkun Lu, Nanxin You).

**Keywords:** Single-cell genomics, Perturbation prediction, Foundation models, Knowledge graphs, Drug discovery, Virtual cells

## Abstract

Understanding cellular responses to genetic perturbations is fundamental for drug discovery, yet experimental approaches face significant limitations in coverage and cost that prevent comprehensive mapping of cellular behavior. This has motivated the development of virtual cells—computational models that learn the relationship between cell state and function to predict the consequences of perturbations across diverse contexts. However, current computational methods suffer from limited accuracy in complex genetic interactions, poor biological interpretability, and inadequate generalization to unseen genes, severely constraining virtual cell capabilities. We present scPert, a multi-modal framework based on Transformer architecture that integrates large language model embeddings with structured biological knowledge to predict single-cell transcriptomic responses to genetic perturbations. Through hierarchical fusion of knowledge graph representations, contextual embeddings from foundation models, and gene-specific encodings, scPert achieves significant performance improvements in both single-gene and combinatorial perturbations over existing methods. In cancer-relevant applications, scPert demonstrates the capability to reveal p53 pathway dynamics and immune checkpoint regulatory mechanisms. Systematic evaluation on 42 cancer dependency genes demonstrates scPert’s ability to identify critical potential therapeutic targets. Our framework establishes a powerful computational foundation for virtual cell construction and accelerates drug target discovery.

## 1. INTRODUCTION

Drug discovery faces the severe challenges of high costs and lengthy development cycles^1–3^, with one of the primary bottlenecks being the lack of systematic understanding of drug mechanisms at the cellular level^4–6^. With the rapid development of single-cell sequencing technologies^7–9^, we can now observe cellular changes under drug or genetic perturbations with unprecedented resolution, opening new possibilities for understanding cellular response mechanisms^4, 10, 11^. However, single-cell perturbation experiments are expensive and yield sparse data that cannot cover all gene combinations or drug treatment conditions^12–14^, making the development of high-precision perturbation effect prediction algorithms crucial for overcoming these limitations^15^. Accurately predicting single-cell transcriptomic changes following genetic perturbations not only helps us understand gene functions and regulatory networks^16^, but more importantly, lays the foundation for future construction of virtual cell systems^5^. Such systems hold the promise of predicting cellular expression profile changes under arbitrary genetic perturbations or drug treatments^17^, thereby greatly accelerating drug screening and combination strategy exploration^15^, and providing powerful computational support for precision medicine and personalized therapy^18^.

Current gene perturbation effect prediction methods can be broadly categorized into several main approaches: foundation model-based methods, generative model-based methods, knowledge-guided methods, and hybrid methods. Foundation model-based methods leverage large-scale pre-trained models on single-cell transcriptomic data, including transformer-based architectures such as scGPT^19^, scFoundation^20^, and GeneCompass^21^. These methods learn latent patterns through self-supervised pre-training on massive datasets and can be fine-tuned for perturbation prediction^22^. A related development involves incorporating language model embeddings from foundation models trained on gene sequences or biological text, exemplified by scGenePT^23^, GenePert^24^, and Scouter^25^, which utilize pre-trained embeddings from models like GenePT^26^ to enhance perturbation prediction. Generative model-based methods, such as Squidiff^27^, employ diffusion processes to model cellular state transitions and predict perturbation responses through probabilistic generation. Knowledge-guided methods explicitly integrate structured biological knowledge, such as gene regulatory networks^28^, protein-protein interaction networks^29^, and metabolic pathway information^30^, into prediction models through graph neural networks^31^.Methods such as GEARS^14^, which leverages gene interaction priors, and BioDSNN^32^, which incorporates pathway constraints, demonstrate how domain knowledge can enhance both interpretability and predictive performance. Additionally, methods like CPA^15^ and biolord^33^ use compositional and disentanglement strategies to improve generalization, while CellOT^34^ applies optimal transport theory to model perturbation responses. These diverse approaches have collectively advanced our ability to predict cellular responses to genetic perturbations, with foundation models excelling at capturing complex patterns from large-scale data^35^ and knowledge-guided methods providing mechanistic interpretability.

However, each approach faces fundamental limitations that constrain their practical utility in drug discovery and systems biology. Foundation model-based methods often function as “black boxes” lacking biological interpretability^35^, making it difficult to derive mechanistic insights for therapeutic target identification. Recent work has shown that even simple linear models can achieve competitive performance with complex deep learning approaches^36^, raising questions about whether current methods truly capture underlying biological mechanisms or merely fit statistical correlations. Methods relying solely on sequence-based or text-based language model embeddings^25, 26^ may not fully exploit the rich structured knowledge in biological databases, such as hierarchical pathway relationships, protein interaction networks, and gene functional annotations, while purely knowledge-guided approaches are constrained by incomplete or inaccurate prior knowledge^32^, particularly for poorly characterized genes. Furthermore, most methods struggle with generalization to unseen genes, novel perturbation combinations, or new cell types^35^, which is critical for drug discovery where therapeutic targets often involve understudied genes or rare cellular contexts^37^. The integration of multi-modal biological information remains largely unexplored, representing a promising but underutilized direction for improving both prediction accuracy and biological interpretability.

To address these challenges, we propose a novel multi-modal fusion Transformer framework for predicting single-cell transcriptomic expression changes following genetic perturbations. Our method employs a hierarchical embedding fusion strategy that integrating knowledge graph-based gene structural information from knowledge graphs with scGPT-derived functional representations from scGPT pre-trained models^19, 38^. We design specialized gene interaction layers and self-attention mechanisms to capture complex regulatory relationships and long-range dependencies in gene networks. Additionally, we develop a novel composite loss function that optimizes for both expression magnitude and directionality, proving particularly valuable for predicting gene expression directionality and identifying key regulatory genes. To address the computational complexity arising from such interactions, we also introduce memory-efficient attention mechanisms and perturbation fusion modules to handle large-scale gene sets and multi-gene combination perturbations. Through comprehensive evaluation on benchmark datasets, our algorithm significantly outperforms existing methods in prediction accuracy and generalization ability, providing a powerful tool for single-cell gene perturbation effect prediction that facilitates the discovery of key regulatory genes and holds significant promise for drug discovery and virtual cell construction.

## 2. Results

### 2.1 LLM-Knowledge Hybrid Framework for Gene Expression Perturbation Prediction

Single-cell perturbation prediction remains a fundamental challenge in computational biology, with recent foundation models showing promise but critical limitations. Geneformer^39^ pioneered the application of transformer architectures to genomics as the first foundation model in this domain, while subsequent single-cell foundation models like scGPT^19^ and scFoundation^20^ further advanced the field. However, these approaches primarily focus on individual cell state modeling without explicitly incorporating perturbation-specific biological knowledge^40^. Knowledge-based methods such as GEARS^14^ and BioDSNN^32^ have demonstrated the value of structured biological relationships but lack the contextual understanding capabilities of large language models.

We developed scPert, a hybrid framework that synergistically combines large language model embeddings with structured biological knowledge to predict gene expression changes following cellular perturbations (**Fig. 1**). Our approach integrates three complementary embedding strategies: (*a*) knowledge graph-based perturbation representations, (*b*) LLM-derived contextual embeddings, and (*c*) gene-specific encodings. This multi-modal fusion enables the model to leverage both the semantic understanding of language models and the precision of curated biological relationships. The framework demonstrates superior performance across diverse perturbation scenarios by explicitly modeling the relationship between perturbation context and cellular response. Unlike previous methods that treat perturbations as discrete entities, scPert captures the semantic similarity between related perturbations, enabling more accurate predictions for previously unseen perturbation combinations and cell types.

**Figure 1.**
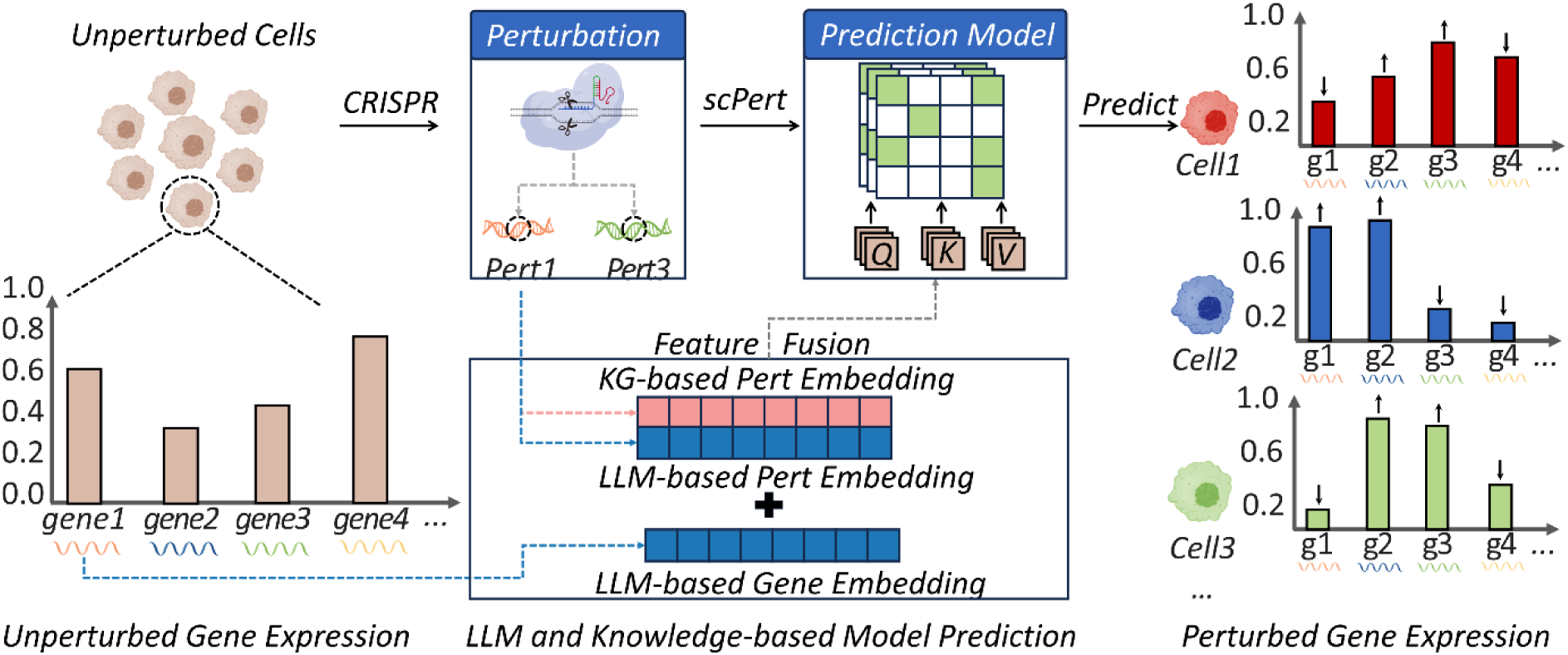
Overview of scPert approach for single-cell perturbation prediction. scPert predicts gene expression changes in single cells following genetic perturbations. The method integrates multi-modal embeddings including knowledge graph-based perturbation embedding, LLM-based perturbation embedding, and LLM-based gene embedding. Through feature fusion and a prediction model, scPert takes gene and perturbation information as input to predict post-perturbation gene expression changes across individual cells, enabling computational prediction of cellular responses to genetic interventions.

### 2.2 Multi-modal architecture enables comprehensive perturbation modeling

#### 2.2.1 Dual-embedding architecture enables comprehensive perturbation modeling

The scPert architecture incorporates several key innovations that distinguish it from existing approaches (**Fig. 2a, b**). We integrate multi-modal embeddings that combine structured biological knowledge from knowledge graphs with data-driven representations from foundation models. This dual approach enables the model to leverage both established biological understanding and patterns learned from large-scale genomics data, addressing a fundamental limitation of purely knowledge-based methods or purely data-driven approaches.

**Figure 2.**
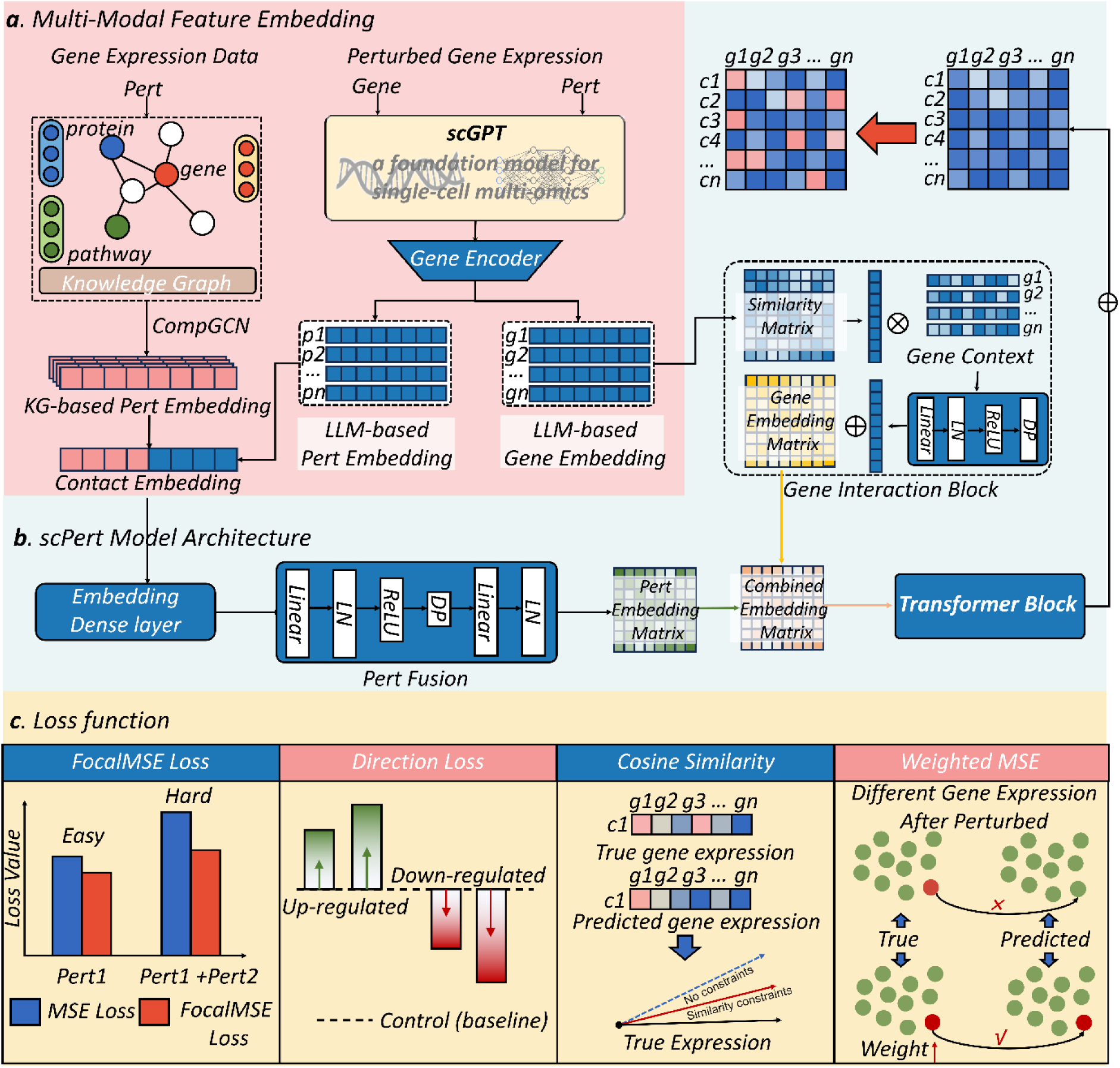
scPert model architecture and multi-objective loss function. (*a*) scPert integrates two complementary embedding strategies: knowledge graph-based perturbation embedding from CompGCN using protein, pathway and other biological networks, and LLM-based perturbation and gene embeddings from the scGPT foundation model. The framework processes gene expression data to generate comprehensive representations for downstream prediction. (***b***) scPert model architecture. The transformer-based architecture employs an embedding dense layer for perturbation fusion, processes gene context through similarity matrices and gene interaction blocks, and uses transformer blocks to model gene-perturbation interactions through cross-attention mechanisms. The model handles both single and combinatorial perturbations through specialized fusion strategies. (***c***) Multi-objective loss function. The training employs a composite loss combining FocalMSE loss (adaptively weighting prediction errors based on sample difficulty, where the x-axis represents perturbation conditions and y-axis shows loss values), Direction loss (enforcing correct gene regulation patterns for up-regulated and down-regulated genes), Cosine Similarity loss (preserving global expression signatures between true and predicted gene expression), and Weighted MSE (accounting for different gene expression patterns after perturbation). This multi-objective optimization ensures biological plausibility while maintaining numerical accuracy. All panels show schematic representations illustrating the operational principles.

Our memory-efficient transformer design handles the computational demands of modeling interactions between thousands of genes and perturbations while maintaining scalability. Critically, we implement specialized mechanisms for combinatorial perturbations that explicitly model non-additive effects—a biological reality that previous methods often oversimplify^15^. The hierarchical gene interaction modeling first captures baseline co-expression patterns before integrating perturbation-specific effects, ensuring that predictions respect both steady-state cellular organization and dynamic perturbation responses.

#### 2.2.2 Multi-objective optimization ensures biological plausibility

A key innovation lies in our multi-objective loss function that addresses multiple aspects of perturbation response prediction simultaneously (**Fig. 2c**). The FocalMSE loss adaptively weights prediction errors based on sample difficulty, proving particularly effective for complex multi-gene perturbations. Our direction loss explicitly enforces correct gene regulation patterns—a critical biological constraint that previous methods often neglect. The cosine similarity component preserves global expression signatures, while weighted MSE accounts for gene-specific heterogeneity.

This comprehensive optimization strategy ensures that predictions maintain biological plausibility while achieving superior numerical accuracy. The model successfully predicts both fine-grained gene-specific changes, establishing a new paradigm that combines the contextual understanding of foundation models with the precision of knowledge-based approaches like GEARS and BioDSNN.

### 2.3 scPert demonstrates superior performance across diverse experimental contexts

To evaluate the generalization capability of our scPert framework, we conducted comprehensive benchmarking using large-scale single-cell perturbation datasets spanning multiple cell types and experimental conditions (**Fig. 3a**). Our evaluation encompassed datasets ranging from smaller focused screens (20-87 perturbations; 44,735-68,603 cells) to large-scale genome-wide screens (up to 1,543 perturbations; over 162,751 cells) (**Note S1, Fig. S1-S2, Table S1**) ^4, 16, 41, 42^, spanning diverse experimental designs and biological contexts. To ensure rigorous evaluation of generalization capabilities, we employed a held-out gene strategy where models were trained without exposure to specific target genes, then tested on predicting the effects of perturbing these previously unseen genes.

**Figure 3.**
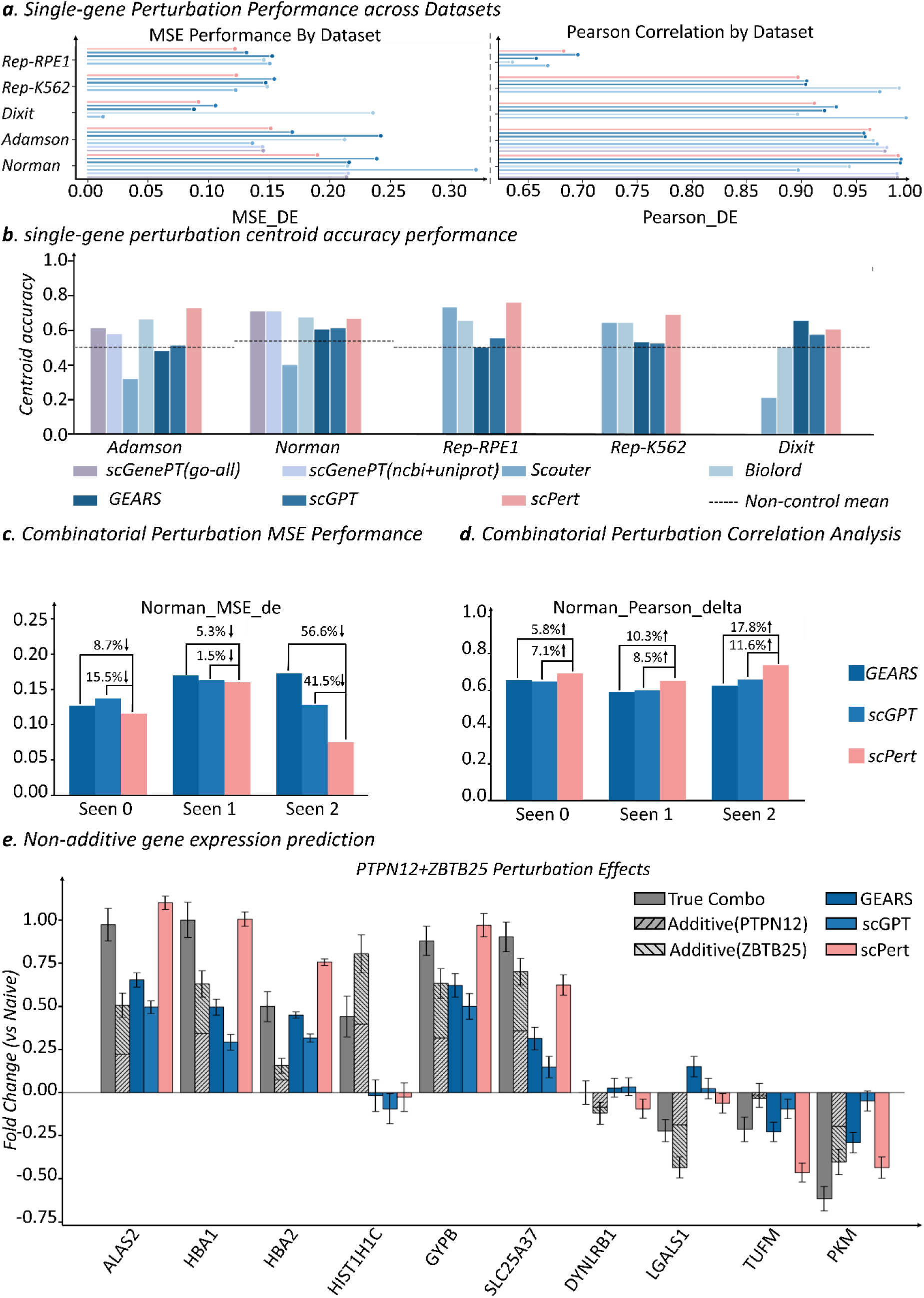
scPert demonstrates superior performance in single-gene and combinatorial perturbation prediction across diverse experimental contexts. (*a*) Performance comparison of scPert against baseline methods across five single-cell perturbation datasets. Left panel shows mean squared error for differentially expressed genes (MSE_DE), and right panel shows Pearson correlation for differentially expressed genes (Pearson_DE) across five benchmark datasets (Norman, Adamson, Dixit, Replogle K562, and Replogle RPE1). scPert consistently achieves competitive prediction errors and maintains high correlations across datasets ranging from small-scale focused screens to large-scale genome-wide studies. (*b*) Single-gene perturbation centroid accuracy performance. Centroid accuracy measures whether predicted profiles are closer to their correct perturbation centroid than to other perturbations’ centroids, with values of 1 indicating perfect recovery of transcriptional effects. The dashed line represents the non-control mean baseline. scPert demonstrates competitive or superior performance across all datasets, particularly excelling in the Dixit dataset. (*c*) Combinatorial perturbation prediction performance using the Norman dataset with 131 gene pairs. Performance is evaluated across three generalization scenarios: “Seen 0” (neither gene observed individually during training), “Seen 1” (one gene previously seen), and “Seen 2” (both genes previously seen individually). scPert shows substantial improvements, particularly in the “Seen 2” scenario with 56.6% reduction in MSE_DE compared to GEARS. (*d*) Pearson correlation delta analysis for combinatorial perturbations, measuring perturbation-induced changes in gene co-expression patterns. scPert demonstrates superior preservation of gene regulatory network structure across all scenarios, with the most pronounced improvements in “Seen 1” and “Seen 2” conditions. (*e*) Prediction accuracy for the top ten non-additively expressed genes in response to PTPN12 and ZBTB25 dual perturbation. scPert accurately captures non-linear combinatorial effects, correctly predicting both synergistic upregulation and suppressive downregulation patterns that cannot be predicted from simple linear addition of individual perturbation effects. Error bars represent standard deviation across multiple evaluations.

We compared scPert against five established baseline methods (**Note S2**): GEARS, a knowledge graph-enhanced approach that represents the current state-of-the-art in perturbation prediction; scGPT, a foundation model specifically designed for single-cell genomics; scGenePT^23^, which augments scGPT with additional gene language representations derived from unstructured biological knowledge; Scouter^25^, a method that uses large language model-derived gene embeddings combined with a compressor–generator neural network to predict transcriptional responses; and biolord^33^, a deep generative framework that disentangles single-cell multi-omic representations into known and unknown biological attributes and uses this disentangled latent space to generate counterfactual cellular profiles, including responses to unseen genetic and drug perturbations.

Our evaluation focused on three complementary metrics: MSE (**Note S4**) for differentially expressed genes (MSE_DE), Pearson correlation for differentially expressed genes (Pearson_DE), and centroid accuracy^43^. Centroid accuracy measures whether predicted post-perturbation profiles are closer to their correct ground-truth centroid than to centroids of other perturbations, using the perturbed centroid as reference to reduce correlation with systematic variation. This metric distinguishes “informative predictions” from “uninformative predictions”, with values of 1 indicating perfect recovery of expected transcriptional effects. We specifically focused on differentially expressed genes for MSE_DE and Pearson_DE because the vast majority of genes show minimal variation between unperturbed and perturbed states, making the prediction of truly responsive genes the more challenging and biologically relevant task.

Based on the latest comprehensive benchmarking, scPert consistently demonstrated competitive or superior performance compared with five state-of-the-art baseline models across diverse single-gene perturbation datasets, indicating strong accuracy and robustness across experimental settings (**Fig. 3a, b**). In terms of MSE_DE (**Fig. 3a, Table S2**), scPert ranked among the top-performing methods across all five benchmarks. Notably, scPert achieved the lowest prediction error in the Norman (0.190) and Replogle RPE1 (0.122) datasets. In the Replogle K562 dataset, scPert (0.123) achieved near-optimal performance, closely matching the best-performing method, Scouter (0.122). In the Adamson dataset, scPert (0.151) showed competitive performance, trailing Scouter (0.136) and scGenePT variants (0.144–0.145), while maintaining balanced correlation performance. Even in the challenging Dixit dataset, where Scouter achieved the lowest MSE_DE(0.013) and GEARS also performed strongly (0.088), scPert remained highly competitive with an error of 0.091. For example, in the RPE1 dataset, scPert yielded substantially lower prediction errors than both GEARS and scGPT (0.122 vs. 0.152 and 0.131, respectively), highlighting its robustness across cellular contexts.

Consistent with these findings, scPert also demonstrated strong Pearson correlation performance across datasets (**Table S3**). While not always achieving the highest correlation, scPert maintained consistently high correlations across all five benchmarks, supporting the stability of its predictions across diverse perturbation settings. In contrast, scGenePT results were available only for the Norman and Adamson datasets due to limited checkpoint availability, where it achieved comparable correlation performance but could not be evaluated on the remaining benchmarks.

In centroid accuracy evaluation (**Note S5, Fig. 3b, Table S4**), scPert exhibited particularly robust and consistent performance across all five datasets. In the Dixit dataset, where most baseline methods showed reduced accuracy (ranging from 0.210 to 0.650), scPert achieved a centroid accuracy of 0.600, ranking second only to GEARS (0.650) and substantially outperforming Scouter (0.210) and the non-control mean baseline (0.500). Across the remaining datasets, scPert consistently ranked at or near the top: achieving the highest centroid accuracy in Adamson (0.724) and Replogle K562 (0.754), the best performance in Replogle RPE1 (0.684), and competitive performance in Norman (0.663), comparable to the strongest baseline methods.

To mechanistically understand scPert’s superior centroid accuracy, we conducted ablation experiments removing the Hetionet knowledge graph component (**Table S10**). Results revealed substantial performance degradation across all datasets. Centroid accuracy decreased from 0.724 to 0.557 in Adamson, from 0.663 to 0.591 in Norman, and from 0.754 to 0.522 in Replogle K562. Most strikingly, in Dixit and Replogle RPE1, removing knowledge graph integration reduced performance to 0.500, exactly matching random baseline and indicating complete loss of ability to distinguish perturbation-specific signals from systematic variation. These results demonstrate that structured biological knowledge graphs provide essential constraints that guide the model toward learning genuine regulatory mechanisms rather than fitting systematic technical artifacts, explaining why scPert achieves robust performance across diverse experimental contexts.

Importantly, scPert scaled effectively from small-scale perturbation screens, such as Dixit, to large-scale genome-wide studies, such as the Replogle datasets, underscoring its generalizability across varying perturbation complexities and experimental designs.

### 2.4 scPert excels in predicting complex combinatorial perturbation effects

#### 2.4.1 Exceptional performance in challenging combinatorial scenarios

Combinatorial genetic perturbations represent a fundamental biological reality where gene function emerges through complex regulatory networks rather than isolated pathways^44^. In cellular systems, most phenotypes result from the interplay of multiple genes, with combinatorial effects often exhibiting non-additive interactions including synergy, epistasis, and buffering mechanisms that cannot be predicted from individual gene perturbations alone^41^. Understanding these combinatorial effects is crucial for therapeutic development, as most diseases involve dysregulation of multiple pathways^45^, and effective treatments often require targeting gene combinations to overcome redundancy and resistance mechanisms inherent in biological systems^46^.

To evaluate scPert’s capability for predicting these complex genetic interactions, we assessed its performance on two-gene combinatorial perturbations using the Norman dataset containing 131 gene pairs. We categorized perturbation combinations into three generalization scenarios based on training data exposure: “Seen 0” where neither gene in the combination had been individually observed during training, “Seen 1” where only one gene was previously seen perturbed individually, and “Seen 2” where both genes had been individually perturbed in the training data, representing progressively easier prediction tasks that mirror real-world drug discovery scenarios where novel combinations must be evaluated (**Fig. 3c, d**).

scPert demonstrated exceptional performance across all combinatorial scenarios, achieving substantial improvements over existing methods (**Fig. 3c, Table S5**). Most strikingly, in the “Seen 2” scenario where both genes were familiar, scPert achieved a remarkable 56.6% improvement in MSE_DE compared to the GEARS, reducing prediction error from 0.180 to 0.078. This dramatic improvement suggests that scPert’s architecture is particularly well-suited for generalizing from observed perturbations to entirely novel gene combinations. In the intermediate “Seen 1” scenario where only one gene had training examples, scPert maintained a slight 5.3% improvement, while in the “Seen 0” baseline condition where neither gene was previously seen individually, it still achieved a modest but consistent 8.7% enhancement over competing methods.

The correlation analysis further reinforced superior performance of scPert in capturing gene expression relationships during combinatorial perturbations (**Fig. 3d, Table S6**). To better understand how perturbations reshape gene regulatory networks, we shifted our focus from static correlation measures (Pearson_DE) to the Pearson correlation delta metric, which specifically quantifies perturbation-induced changes in gene co-expression patterns (**Note S3**). scPert showed the most pronounced improvements in common scenarios, with substantial enhancements in Pearson correlation delta for “Seen 1” and “Seen 2” conditions, respectively. This preservation of correlation structures provides valuable insights into how combinatorial perturbations reshape gene regulatory networks. Such capabilities may contribute to our understanding of complex cellular responses to multi-target interventions.

The progressive difficulty across scenarios reveals important insights about cellular robustness and genetic interaction landscapes. The “Seen 2” scenario represents the easiest condition where both genes have individual training examples, while “Seen 0” poses the greatest challenge by requiring prediction of entirely novel gene combinations that were never observed individually during training. scPert’s robust performance across this difficulty gradient, particularly its superior correlation preservation in the most challenging “Seen 0” scenario, demonstrates its ability to leverage learned representations of gene function and regulatory network topology to extrapolate to novel combinations. This capability addresses a critical bottleneck in systems biology where the combinatorial space of possible perturbations vastly exceeds experimental feasibility, making computational prediction essential for prioritizing experiments and understanding genetic interaction landscapes.

#### 2.4.2 Biological insights from non-linear perturbation effects

To illustrate scPert’s practical utility in combinatorial perturbation prediction, we first examined the model’s fundamental capacity to identify key responsive genes. scPert demonstrated robust accuracy in predicting the top 20 differentially expressed genes for both single and combination perturbations (**Fig. S3**), establishing its reliability for downstream analysis of complex genetic interactions. Based on this established predictive accuracy, we investigated why genetic interactions cannot be accurately captured through simple linear addition of individual perturbation effects^16^. Combinatorial perturbations involve complex regulatory crosstalk, feedback mechanisms, and emergent network properties that only manifest when multiple genes are simultaneously perturbed, leading to non-additive transcriptional responses that deviate substantially from the sum of individual effects.

Based on previous work (GEARS^14^), we selected the top ten non-additively expressed genes in response to *PTPN12* and *ZBTB25* dual perturbation for prediction analysis (**Fig. 3e**). scPert predicted the correct non-additive effects across almost all of these top ten non-additively expressed genes following the perturbation of *PTPN12* and *ZBTB25*.

scPert accurately captured the nuanced expression patterns of genes showing non-additive responses, correctly predicting both synergistic upregulation in hemoglobin components (*HBA*^47^) and suppressive downregulation in metabolic genes (*PKM*^48^). The model’s success in identifying cases where modest individual perturbation effects combine to produce dramatic transcriptional changes suggests that combinatorial perturbations involve complex regulatory interactions that extend beyond simple additive effects. This capability positions scPert as a powerful tool for dissecting genetic interaction networks and identifying therapeutic vulnerabilities that emerge specifically from multi-target interventions.

### 2.5 Comprehensive Genetic Interaction Prediction and Analysis

#### 2.5.1 Evaluation of Genetic Interaction Score Predictions

To comprehensively evaluate scPert’s capacity for deciphering complex genetic interaction networks, we conducted systematic prediction analysis across five distinct genetic interaction (GI) modalities: synergy, epistasis, redundancy, suppression, and neomorphism^41^ (**Note S6**). These GI scores quantify different aspects of non-additive genetic interactions, providing a multifaceted view of combinatorial perturbation effects.

We compared scPert against two established methods (GEARS and scGPT) across 125 pairwise gene combinations. For synergistic interactions, scPert demonstrated superior performance with scores ranging from 0.5 to 1.8, closely matching the experimental distribution (**Fig. 4a**). The method showed particularly strong agreement with actual data in capturing high-magnitude synergistic effects, which are critical for identifying functionally cooperative gene pairs.

**Figure 4.**
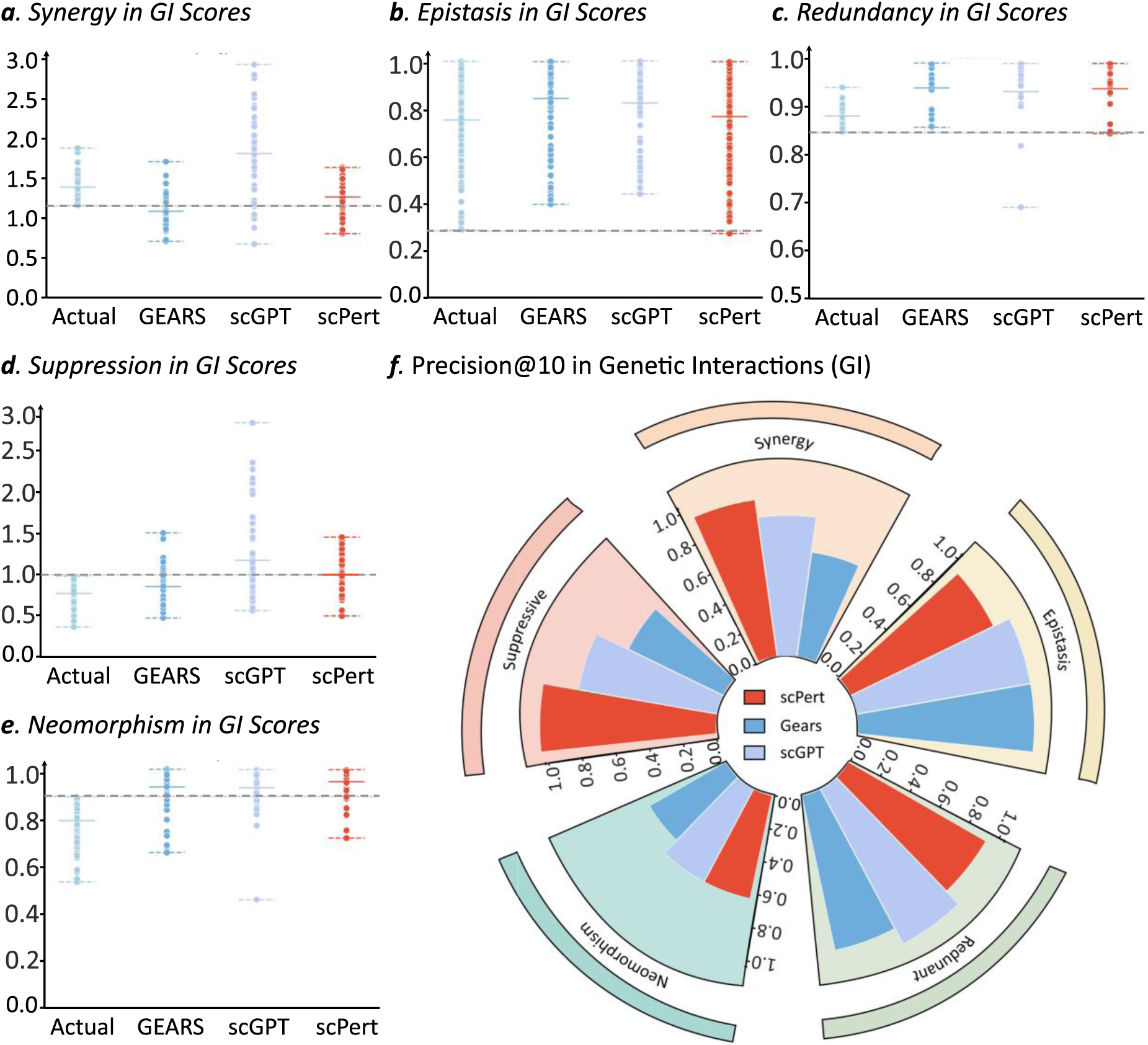
Comprehensive genetic interaction prediction and analysis by scPert. (***a-e***) Genetic interaction score evaluation across five modalities. scPert was compared against GEARS and scGPT across 125 pairwise gene combinations for five genetic interaction types: (*a*) synergy, (*b*) epistasis, (*c*) redundancy, (*d*) suppression, and (*e*) neomorphism. Each plot shows the distribution of predicted GI scores compared to experimental values (Actual). scPert demonstrates superior performance in capturing synergistic interactions and maintains competitive performance across other GI modalities. (***f***) Precision@10 analysis for genetic interaction prediction. The circular plot displays precision@10 metrics measuring the fraction of correctly predicted interactions within the top 10 predictions for each GI subtype (**Table S7**). scPert (red segments) shows notable performance improvements over GEARS (blue segments) and scGPT (light blue segments) across multiple interaction types, with particularly strong performance in synergy and redundancy predictions. This analysis demonstrates scPert’s practical utility for prioritizing genetic interactions for experimental validation.

In epistatic interactions, all computational methods showed similar performance, with scores predominantly clustering around 0.24-1.00 (**Fig. 4b**). This consistency suggests that epistatic relationships, where one gene masks the effect of another, are relatively well-captured by current perturbation prediction approaches. For redundancy scores, scPert maintained high concordance with experimental data, with most predictions falling within the 0.84-1.00 range, indicating effective capture of functionally overlapping gene relationships (**Fig. 4c**).

Suppression interactions revealed more challenging prediction scenarios, with scPert showing moderate performance compared to experimental data (**Fig. 4d**). Notably, scGPT, as a foundation model trained on diverse genomic data, exhibited highly dispersed predictions across all GI types, reflecting the inherent challenge that large-scale pre-trained models face when adapting to specific combinatorial perturbation tasks. This dispersed behavior suggests that while foundation models excel at capturing general genomic patterns, they may lack the specialized architecture needed for precise genetic interaction modeling, particularly for complex multi-gene perturbations where intricate gene-gene relationships must be accurately quantified.

Similarly, neomorphism—representing novel phenotypes emerging from gene combinations—showed variable prediction accuracy across all methods, with scPert achieving reasonable performance while scGPT continued to show the characteristic broad distribution pattern observed across other GI types (**Fig. 4e**). This pattern underscores a fundamental limitation of foundation model approaches in genetic interaction prediction: their general-purpose training may not provide the focused inductive biases necessary for modeling the nuanced, context-dependent nature of combinatorial gene perturbations.

#### 2.5.2 Precision Analysis and Comprehensive Genetic Interaction Mapping

To evaluate the practical utility of our predictions, we assessed the precision@10 metric for each GI type (**Note S7, S8, S9**), measuring how many of the top 10 predicted interactions were experimentally validated. scPert achieved notable precision across different interaction types, with particularly strong performance in synergy and redundancy predictions (**Fig. 4f**). This analysis reveals that scPert reliably identifies the most significant genetic interactions, making it valuable for prioritizing experimental validation.

Building on the validated predictive performance demonstrated in Figure 4, where scPert achieved strong precision across genetic interaction modalities (Precision@10: synergy 0.9, suppressive 1.0), we performed genetic interaction network analysis by systematically predicting interactions among all possible pairwise combinations of 105 genes, generating a comprehensive functional genomics landscape of 5,460 gene pairs (**Fig. 5**). This systems-level approach enabled construction of a multi-dimensional genetic interaction network that simultaneously captures all five GI modalities, providing unprecedented mechanistic insight into the combinatorial gene regulation space.

**Figure 5.**
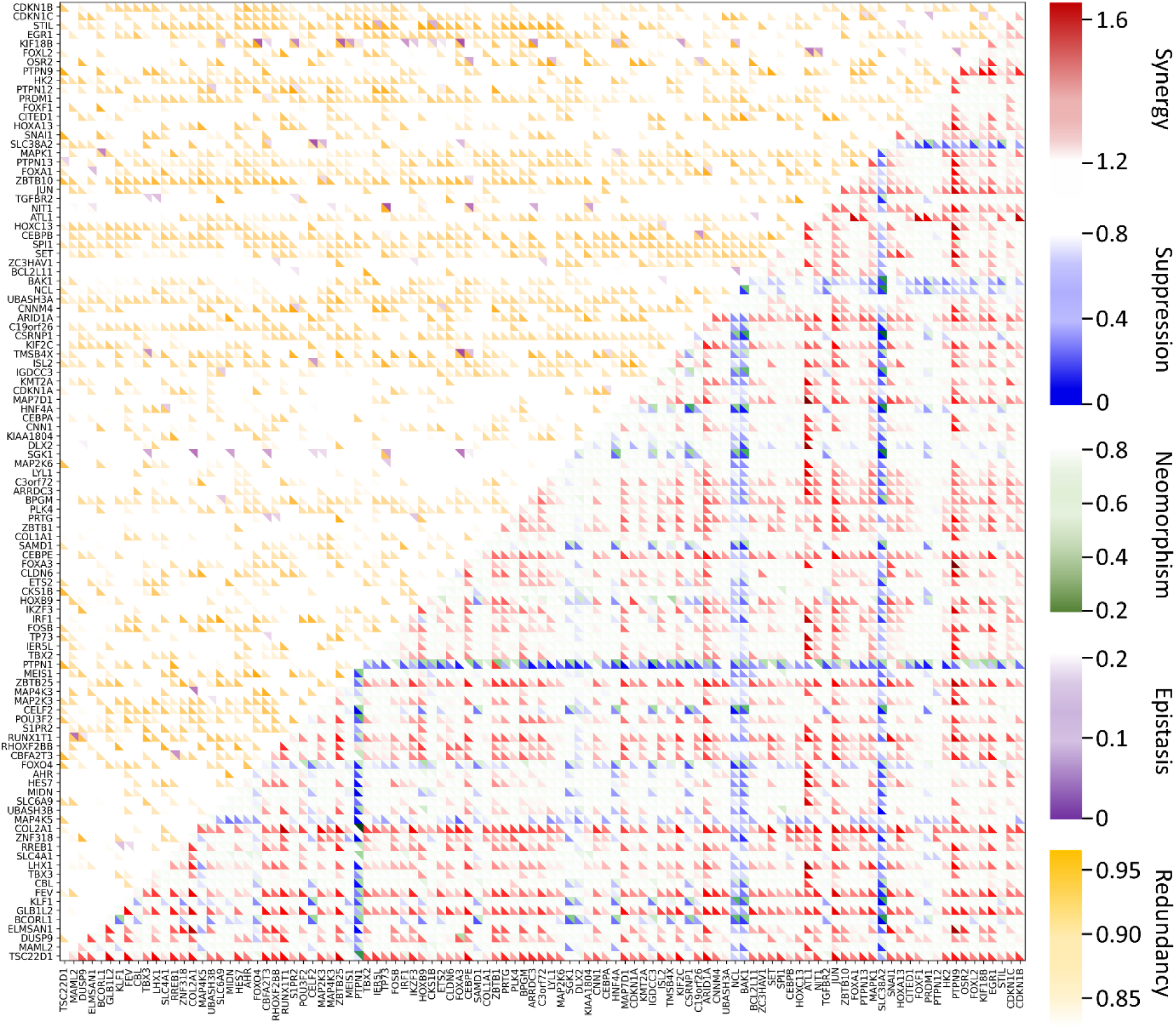
Multidimensional genetic interaction network map for pairwise gene combinations. This multidimensional genetic interaction map displays predicted interactions among all possible pairwise combinations of 105 genes, generating a comprehensive functional genomics landscape of 5,460 gene pairs. Each cell in the matrix represents a gene pair, with different colored triangular segments indicating the strength and type of genetic interaction: synergy (red), suppression (blue), neomorphism (green), redundancy (yellow), and epistasis (purple). The map reveals distinct patterns where functionally related genes exhibit coherent interaction architectures. Genes within the same regulatory pathways often display strong synergistic interactions (red triangles), while genes in compensatory pathways frequently show redundant (yellow triangles) or epistatic (purple triangles) relationships. This systems-level analysis demonstrates scPert’s ability to construct comprehensive functional genomics landscapes that simultaneously capture multiple interaction modalities, providing unprecedented mechanistic insight into the combinatorial gene regulation space and identifying key genetic interaction hubs for potential therapeutic targeting.

The resulting genetic interaction network revealed distinct topological patterns and functional modules, with genes exhibiting pathway-specific interaction preferences. The network analysis demonstrates clear modality-specific perturbational signatures, where genes participating in related biological processes exhibit coherent interaction architectures. The genetic interaction network reveals that genes with different regulatory roles exhibit distinct interaction patterns. For example, structural maintenance genes like *ATL1* tend to show synergistic interactions^49^, while signaling inhibitors such as *PTPN1* display suppressive interactions^50^, reflecting their distinct cellular roles and highlighting the potential of interaction profiles for functional annotation. To further validate the biological fidelity of the inferred interactions, we benchmarked the predicted GI scores against combinatorial cell fitness screen data. Consistent with the validation framework used in GEARS, scPert-predicted scores for experimentally confirmed strong synergistic pairs were significantly distinguishable from those of neutral pairs (P < 0.01, one-sided t-test), achieving a directional consistency of 58.5% across the broad 105-gene landscape, confirming that the network topology reflects genuine biological signals.

This systems-level genetic interaction network analysis represents a paradigm shift from conventional pairwise interaction studies toward comprehensive functional genomics mapping. By simultaneously modeling multiple GI dimensions in a network context, scPert enables researchers to dissect the complex regulatory logic underlying cellular phenotypes and identify key genetic interaction hubs that could serve as therapeutic targets. The capacity of scPert to predict diverse interaction modalities with high fidelity makes it an invaluable computational tool for advancing our understanding of genetic interaction networks and their roles in cellular function and disease.

### 2.6 scPert demonstrates superior accuracy in cancer-relevant gene prediction

#### 2.6.1 Enhanced prediction of p53 pathway perturbations reveals tumor suppressor network dynamics

The p53 tumor suppressor pathway represents one of the most critical regulatory networks in cancer biology, with p53 often referred to as the “guardian of the genome” due to its central role in maintaining genomic stability and preventing malignant transformation. Given that p53 mutations occur in over 50% of human cancers, accurate prediction of p53 pathway perturbations is essential for understanding tumor suppressor mechanisms and developing targeted therapeutic strategies^51^.

To evaluate scPert’s performance on this clinically relevant pathway, we identified 200 p53-related genes from the Reactome database^52^. By intersecting these pathway genes with the 105 single-gene perturbations available in the Norman dataset, we identified five genes with perturbation data: *CEBPA*, *BAK1*, *TSC22D1*, *CDKN1A*, and *JUN*. These genes represent diverse functional components of the p53 network, including transcriptional regulators (*CEBPA*^53^*, JUN*^54^), cell cycle checkpoint proteins (*CDKN1A*^55^), pro-apoptotic factors (*BAK1*^56^), and stress response mediators (*TSC22D1*^57^).

We comprehensively evaluated the three methods—GEARS, scGPT, and scPert—across multiple performance metrics for predicting the top 20 differentially expressed genes following perturbation of these p53 pathway components. scPert demonstrated superior performance across all evaluated metrics (**Fig. 6a, Table S8**), achieving the highest Pearson correlation (0.927), competitive direction accuracy (0.79), and the lowest mean absolute error (MAE) of 0.153. In comparison, GEARS obtained correlation, direction accuracy, and MAE values of 0.899, 0.69, and 0.169, respectively, while scGPT achieved 0.907, 0.82, and 0.191 for the corresponding metrics (**Note S10**).

**Figure 6.**
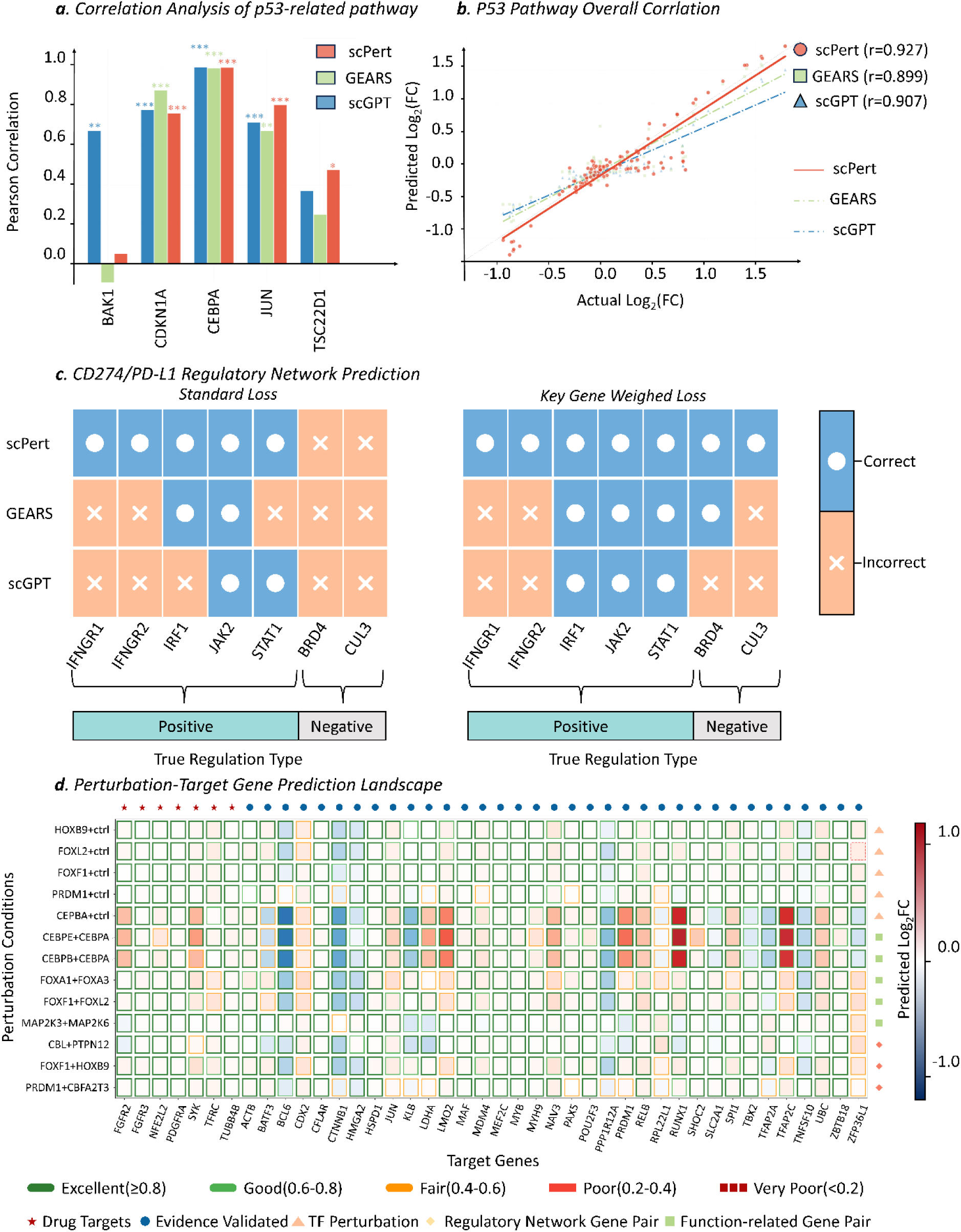
scPert captures biologically meaningful cancer regulatory networks. (***a***) p53 pathway perturbation prediction performance. Pearson correlation analysis for five key p53 network components (CEBPA, BAK1, TSC22D1, CDKN1A, JUN) shows scPert achieving significant correlations (***p < 0.001) for four out of five perturbations, with three genes showing correlations > 0.8. Error bars represent 95% confidence intervals. (***b***) Overall p53 pathway correlation analysis. Scatter plot comparing predicted versus actual log₂ fold changes across p53 pathway genes. scPert demonstrates overall correlation (r = 0.927) compared to GEARS (r = 0.899) and scGPT (r = 0.907), indicating robust capture of p53 pathway-associated transcriptional responses and tumor suppressor network dynamics. (***c***) CD274/PD-L1 regulatory network prediction. Comparison of directional prediction accuracy for PD-L1 regulators using standard loss (left) versus key gene weighted loss (right). Standard optimization shows systematic bias toward positive regulation across all methods. After implementing targeted weighting for CD274 expression, scPert achieves perfect directional accuracy (7/7 correct), correctly identifying both positive regulators (IFNGR1, IFNGR2, IRF1, JAK2, STAT1 - green) and negative regulators (BRD4, CUL3-gray), revealing clinically relevant immune checkpoint regulatory mechanisms. Blue indicates correct predictions, orange indicates incorrect predictions. (***d***) Cancer therapeutic target prediction landscape. Systematic evaluation of 42 cancer-relevant target genes across 13 perturbation conditions, yielding 546 individual predictions. Each cell represents predicted log₂ fold change with colors indicating magnitude and direction (blue: downregulation, red: upregulation). Prediction quality is indicated by border styles: thick green borders (excellent, ≥0.8), thin red borders (poor, <0.2). Gene symbols mark known drug targets (★) and literature-validated genes (●). Results show that 76.9% of predictions achieve excellent quality scores, enabling identification of therapeutic vulnerabilities and drug target prioritization.

The gene-level correlation analysis revealed scPert’s exceptional ability to capture pathway-specific expression signatures (**Fig. 6b**). Among the five p53 pathway perturbations, scPert achieved statistically significant correlations (p < 0.001) for four perturbations (CDKN1A, JUN, CEBPA with r > 0.8), while GEARS showed significant correlations for only three perturbations and scGPT for four. Notably, scPert’s overall correlation of r = 0.927 indicates robust capture of p53-mediated transcriptional responses, suggesting its potential utility for predicting therapeutic targets within tumor suppressor networks.

#### 2.6.2 Accurate prediction of immune checkpoint regulation in limited data scenarios

Immune checkpoint inhibitors have revolutionized cancer treatment, with PD-L1 (encoded by *CD274*) serving as a key therapeutic target^58^. Understanding the regulatory networks controlling PD-L1 expression is crucial for optimizing immunotherapy efficacy and identifying resistance mechanisms. Previous work by Papalexi et al.^59^ identified key regulators of PD-L1 expression, including positive regulators (*IFNGR1, IFNGR2, IRF1, JAK2, STAT1*) and negative regulators (*BRD4, CUL3*), using ECCITE-seq technology^60^.

To evaluate scPert’s performance in clinically relevant small-data scenarios, we applied our framework to the same scCRISPR-seq dataset containing 99 perturbations in THP-1 cells (human monocytic leukemia line). After quality filtering (**Note S11**), we identified 11 effective perturbations for analysis. This dataset presents unique challenges due to its limited size and the biological complexity of immune checkpoint regulation in cancer cells.

Initial predictions revealed a systematic bias toward positive regulation across all methods, with scPert correctly predicting 5 out of 7 regulatory directions (all positive regulators: *IFNGR1, IFNGR2, IRF1, JAK2, STAT1*), while GEARS and scGPT each correctly predicted only 2 out of 7 directions. This limitation likely reflects two key factors: (a) dataset imbalance, with only 2 known negative regulators among 11 perturbations, leading to insufficient training examples for negative regulation patterns; and (b) the dilution effect of *CD274* expression signals among 1,800+ genes during standard MSE optimization, causing the model to underweight this specific target gene.

To address these limitations, we implemented a targeted weighting strategy that enhanced the model’s attention to CD274 expression changes. Following this optimization, scPert achieved perfect directional prediction (7/7 correct), accurately identifying both positive and negative PD-L1 regulators. In contrast, GEARS improved to 4/7 correct predictions (3 positive, 1 negative), while scGPT reached 3/7 correct predictions (all positive). This substantial improvement demonstrates scPert’s adaptability to small-scale, clinically focused datasets and its potential for identifying therapeutic targets in precision medicine applications (**Fig. 6c**).

The successful prediction of immune checkpoint regulation highlights scPert’s utility for understanding complex regulatory networks in cancer immunotherapy. The model’s ability to capture both positive and negative regulatory relationships, even with limited training data, suggests its potential for predicting novel therapeutic combinations and resistance mechanisms in checkpoint inhibitor therapy.

#### 2.6.3 Systematic identification of cancer therapeutic targets through perturbation-guided dependency mapping

To evaluate scPert’s performance on clinically relevant cancer targets, we leveraged the second-generation Cancer Dependency Map (DepMap)^61, 62^, a comprehensive resource that systematically identifies dependencies across cancer cell lines and provides a framework for target prioritization. From this clinically informed dependency map, we extracted pan-cancer essential genes categorized into three groups based on therapeutic tractability: targets with approved or pre-clinical drugs (group 1), targets with supporting evidence of future tractability (group 2), and those with weak or no evidence of future tractability (group 3)^63^. We focused on groups 1 and 2, representing the most clinically relevant targets, and intersected these with genes having experimental perturbation data in the Norman dataset, yielding a curated set of 42 cancer-relevant target genes including known drug targets (*FGFR2, FGFR3, NFE2L2, PDGFRA, SYK, TFRC, TUBB4B*) and literature-validated therapeutic candidates.

For perturbation conditions, we systematically selected three types of perturbations from the pool of unseen experimental conditions that the model had not encountered during training: (a) Single transcription factor perturbations: critical regulatory factors with well-defined individual effects, including *CEBPA* (hematopoietic differentiation^64^), *SPI1* (myeloid lineage specification^65^), *PRDM1* (B-cell differentiation and cell cycle control^66^), *ETS2* (ETS family developmental regulator^67^), *FOXF1* (mesenchymal development^68^), *FOXL2* (ovarian development^69^), *HOXB9* (body axis patterning^70^), and *TBX2* (T-box developmental control^71^). (b) Functionally related dual perturbations: gene pair combinations with known synergistic or antagonistic relationships, including transcription factor family cooperations (CEBPA+CEBPB, CEBPA+CEBPE, FOXA1+FOXA3), signaling pathway components (MAP2K3+MAP2K6, CBL+PTPN12), and immune regulators (UBASH3A+UBASH3B). (c) Regulatory network perturbations: transcriptional crosstalk combinations such as TBX2+TBX3, FOXF1+HOXB9, and PRDM1+CBFA2T3.

The final evaluation encompassed 13 perturbation conditions across 42 cancer-relevant target genes, yielding 546 individual predictions (**Fig. 6d**). scPert demonstrated exceptional performance with 76.9% of predictions achieving excellent quality scores (≥0.8), 12.5% reaching good levels (0.6-0.8), and only 10.6% falling below good quality thresholds (**Note S12, Fig. S4**). Critically, the model successfully identified *RUNX1* as a key downstream target consistently regulated across multiple hematopoietic transcription factor perturbations. Additionally, combination perturbations involving *CEBPA* (CEBPA+CEBPB, CEBPA+CEBPE) exhibited remarkably similar expression profiles to *CEBPA* single perturbation across the target gene set, suggesting that CEBPA exerts dominant regulatory effects in these combinatorial contexts (**Fig. S5**).

The biological significance of these computational discoveries is underscored by extensive clinical evidence. *RUNX1* and *CEBPA* function as interconnected transcriptional regulators in acute myeloid leukemia (AML) pathogenesis, both recruiting *TET2* demethylase to chromatin and facilitating DNA demethylation to maintain normal hematopoietic gene expression programs. In AML, mutations in either factor disrupt this regulatory axis, leading to hypermethylation of target gene promoters and transcriptional silencing of tumor suppressors^72^. The t(8;21) translocation, which generates the oncogenic RUNX1-RUNX1T1 fusion, specifically interferes with CEBPA-mediated autoregulation and blocks normal myeloid differentiation programs^73^. The observed CEBPA dominance in combination perturbations further supports its role as a master regulator whose transcriptional program can override co-perturbed factors, positioning it as a high-priority therapeutic target.

Most importantly, scPert’s ability to systematically map these regulatory dependencies from limited experimental data opens new avenues for cancer target discovery. By accurately predicting how perturbations of upstream regulators affect downstream oncogenes and tumor suppressors, the model enables researchers to identify previously uncharacterized therapeutic vulnerabilities and prioritize combination treatment strategies that target interconnected regulatory nodes. This computational approach significantly accelerates the discovery of cancer dependencies that would otherwise require extensive experimental screening, positioning scPert as a powerful tool for precision oncology research.

## 3. Discussion

We introduce scPert, a hybrid computational framework that synergistically combines large language model representations with structured biological knowledge to predict single-cell transcriptomic responses to genetic perturbations. By integrating foundation model capabilities with curated biological relationships, scPert addresses fundamental limitations of existing approaches and establishes a new paradigm for perturbation response prediction.

Our framework introduces several critical innovations that distinguish it from existing approaches. The multi-modal embedding fusion strategy effectively bridges the gap between data-driven foundation models and knowledge-based methods, leveraging both the semantic understanding of transformer architectures and the precision of curated biological relationships. Unlike previous methods that treat perturbations as discrete entities, scPert captures semantic similarity between related perturbations through its hierarchical embedding fusion, enabling more accurate predictions for novel gene combinations and cell types. The specialized architecture for combinatorial perturbations represents a significant advance in modeling complex genetic interactions, successfully predicting emergent regulatory modules and pathway crosstalk that arise specifically from multi-target interventions.

The rapid development of single-cell foundation models has highlighted the power of large-scale pretraining to capture global transcriptional patterns. However, purely data-driven approaches often lack perturbation-specific inductive biases^36, 74^, while knowledge-based methods provide biological interpretability at the cost of flexibility. In addition to hybrid fusion strategies, recent graph-aware transformer models such as GREMLN^75^ incorporate regulatory network topology directly into the attention mechanism, representing a complementary avenue for integrating structured biological knowledge into single-cell modeling. Across benchmarks, we observed consistent performance patterns associated with dataset scale and evaluation metrics. On smaller perturbation screens, some methods achieved strong results on individual metrics, whereas larger genome-wide datasets revealed greater performance divergence. While most approaches maintained high Pearson correlations, stricter metrics such as centroid accuracy exposed substantial variability, indicating that correlation-based evaluations may mask systematic prediction errors. By integrating foundation model representations with structured biological knowledge, scPert balances expressiveness and biological grounding, resulting in robust performance across datasets and evaluation criteria.

The superior performance in cancer-relevant applications demonstrates scPert’s potential for therapeutic target discovery. Our systematic evaluation of p53 pathway perturbations reveals exceptional ability to capture tumor suppressor network dynamics, achieving correlations of r = 0.927 with experimental data. The successful prediction of immune checkpoint regulation, particularly the accurate identification of PD-L1 regulatory networks, highlights the framework’s utility for immunotherapy research and resistance mechanism discovery. The genetic interaction network analysis represents a paradigm shift from conventional pairwise studies toward comprehensive functional genomics mapping, enabling researchers to dissect complex regulatory logic underlying cellular phenotypes and identify key genetic interaction hubs that could serve as potential therapeutic targets.

Genetic interactions have been increasingly recognized as critical determinants of disease phenotypes and therapeutic responses, with systematic mapping across model organisms and human diseases revealing that interactions preferentially bridge distinct compensatory functional modules and harbor targetable vulnerabilities for combination therapies^76–78^. Particularly noteworthy is scPert’s ability to predict diverse genetic interaction types with high precision, including synergistic, epistatic, and suppressive relationships. The model’s success in identifying cases where modest individual perturbation effects combine to produce dramatic transcriptional changes illuminates fundamental principles of cellular regulation, including threshold effects, feedback loops, and compensatory mechanisms that only become apparent through combinatorial analysis. This capability positions scPert as a powerful tool for dissecting genetic interaction networks and identifying therapeutic vulnerabilities that emerge specifically from multi-target interventions.

Despite these advances, several limitations warrant consideration for future development. First, the current framework’s performance on small-scale datasets suggests the need for specialized architectures optimized for limited data scenarios, particularly in clinical applications where sample availability may be constrained. While scPert effectively captures distinct cellular states induced by single perturbations, combination perturbations, and mixed conditions as evidenced by clear clustering patterns and expression distance relationships (**Fig. S6**), the increasing complexity of mixed perturbation scenarios reveals challenges in maintaining prediction accuracy across all experimental contexts. Second, like current perturbation prediction methods, scPert requires dataset-specific training and does not yet achieve cross-dataset generalization without retraining, reflecting a field-wide challenge in developing universal models that can transfer learned perturbation response patterns across diverse cellular contexts and experimental conditions. Third, scPert’s reliance on existing knowledge graph completeness may limit its ability to predict entirely novel biological relationships, indicating opportunities for incorporating causal inference frameworks and dynamic knowledge discovery mechanisms. Fourth, while our multi-modal fusion approach enhances biological interpretability compared to pure foundation model methods, the complex embedding representations still present challenges for mechanistic understanding, highlighting the need for continued development of explainable AI approaches in genomics applications.

The computational efficiency and scalability of scPert also merit consideration. While our memory-efficient transformer design enables handling of large gene sets, the computational requirements for training and inference may limit accessibility for smaller research groups. Future developments should focus on model compression techniques and efficient fine-tuning strategies to democratize access to advanced perturbation prediction capabilities. Additionally, standardization of evaluation metrics and benchmarking procedures across the field would facilitate more robust comparison of methods and accelerate methodological improvements.

These limitations point toward promising directions for future development. Advances in perturbation prediction will increasingly benefit from multi-modal integration and expanded information sources. Cross-omics foundation models that jointly learn from transcriptomic, proteomic, metabolomic, and epigenomic layers will capture regulatory mechanisms spanning molecular scales that single-omics approaches cannot reveal. Future frameworks should incorporate diverse information beyond structured biological knowledge, including protein-level representations, unstructured textual information from scientific literature, and other complementary modalities, to enhance prediction comprehensiveness while addressing current knowledge graph limitations, as such data have been increasingly integrated into well-curated databases^79–81^. Multi-scale modeling that integrates molecular, cellular, tissue, and organismal predictions will be essential for comprehensive virtual cell systems capable of predicting not only transcriptomic responses but also downstream cellular phenotypes and therapeutic outcomes. Such developments will accelerate the translation from computational predictions to experimental validation and clinical applications, advancing the field toward truly predictive virtual cell platforms for drug discovery.

scPert represents a significant advancement in computational biology by successfully integrating foundation model capabilities with structured biological knowledge for single-cell perturbation prediction. Through superior performance across diverse experimental contexts and demonstrated accuracy in clinically relevant applications, scPert establishes a robust framework for virtual cell construction. The ability to predict complex genetic interactions while maintaining biological plausibility enables the development of computational models that can simulate cellular responses to arbitrary perturbations, ultimately facilitating drug target discovery and therapeutic development. As computational approaches become increasingly essential in modern drug discovery, scPert provides a critical foundation for advancing from virtual cell systems toward practical pharmaceutical applications.

## 4. Methods

### 4.1 Model Architecture Overview

Single-cell perturbation response prediction faces significant challenges due to the high dimensionality of gene expression data and the complex, often non-linear nature of cellular regulatory mechanisms. Traditional approaches often fail to capture the intricate relationships between genes and their responses to genetic perturbations, particularly when dealing with combinatorial interventions.

We introduce scPert, a novel deep learning framework designed to predict gene expression responses to genetic perturbations in single-cell data. Given a perturbation dataset 𝒟 = {(𝐠^𝑖^, 𝒫^𝑖^)}^𝑁^, where 𝐠^𝑖^ ∈ ℝ^𝐾^ represents the gene expression vector of cell 𝑖 with 𝐾 genes, and 𝒫^𝑖^ = (𝑃1^𝑖^, ⋯, 𝑃𝑀^𝑖^) is the set of perturbations of size 𝑀 performed on cell 𝑖 (𝑀 = 0 corresponds to an unperturbed cell), scPert learns a function 𝑓 that maps a novel perturbation set 𝒫 to its post-perturbation gene expression outcome 𝐠.

Our architecture incorporates several key innovations that distinguish it from previous approaches. First, we integrate multi-modal embeddings that combine structured biological knowledge from knowledge graphs with data-driven representations from foundation models. This dual approach allows us to leverage both established biological understanding and patterns learned from large-scale genomics data. Second, we implement a memory-efficient transformer architecture that can handle the computational demands of modeling interactions between thousands of genes and perturbations. Third, we design specialized mechanisms for handling combinatorial perturbations, explicitly modeling the non-additive effects that are common in biological systems. Finally, we incorporate hierarchical gene interaction modeling that first captures baseline co-expression patterns before integrating perturbation-specific effects.

The complete pipeline consists of four main components: (a) embedding generation for both genes and perturbations, (b) perturbation encoding that handles both single and combinatorial interventions, (c) cross-attention mechanisms that model gene-perturbation interactions, and (d) expression prediction with residual connections.

### 4.2 Multi-Modal Gene Representation Learning

#### 4.2.1 Knowledge Graph Embeddings

Understanding gene function and relationships requires incorporating structured biological knowledge that has been accumulated over decades of research. To achieve this, we leverage the Hetionet knowledge graph, a comprehensive biomedical knowledge base containing 47,031 nodes across 11 different entity types and over 2.2 million relationships^38^. This graph encompasses not only genes and pathways but also diseases, molecular functions, anatomical structures, and pharmacological information, providing a rich context for understanding gene behavior.

We employ Composition-based Graph Convolutional Networks (CompGCN^82^, **Note S13, Table S9**) to learn gene embeddings that preserve both structural and semantic relationship. The rationale for KGE integration lies in its ability to encode prior biological knowledge that is often missing in purely data-driven approaches. Unlike traditional methods that treat genes as isolated entities, our KGE representations capture evolutionary relationships, pathway memberships, and functional similarities that are crucial for understanding perturbation effects.

*4.2.2 Foundation Model Embeddings and Fusion*

While knowledge graphs provide structured biological understanding, they may not capture the full complexity of gene expression patterns observed in real single-cell data. To address this limitation, we incorporate pre-trained embeddings from scGPT, a foundation model trained on extensive single-cell genomics datasets. These embeddings capture context-dependent expression patterns, cell-type-specific regulatory states, and co-variation relationships that emerge from actual experimental observations.

The integration of these two complementary representation types occurs through a learned fusion mechanism. For individual perturbations, we concatenate the knowledge graph and foundation model embeddings, then apply a multi-layer perceptron to learn optimal combination strategies:

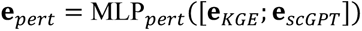

For combinatorial perturbations involving multiple genes, the challenge becomes more complex as we need to model potential interactions between the different perturbations. We address this through a specialized fusion mechanism that averages individual perturbation embeddings before applying a dedicated fusion network:

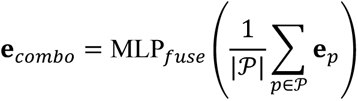

This approach allows the model to learn how different perturbations might interact synergistically or antagonistically, moving beyond simple additive effects.

### 4.3 Gene Interaction and Transformer Architecture

Gene expression is fundamentally a system-level phenomenon where the response of one gene to a perturbation can influence and be influenced by the expression of many other genes. To capture these complex interdependencies, we implement a self-attention mechanism that computes contextual relationships between genes:

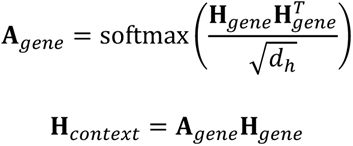

where 𝐇_𝑔𝑒𝑛𝑒_ ∈ ℝ^𝑁×𝑑ℎ^ represents the processed gene embeddings, and the attention mechanism allows genes to attend to one another based on their learned representations. This approach is crucial for capturing regulatory relationships, co-expression patterns, and pathway-level effects that might not be explicitly encoded in the knowledge graph.

We process these interactions through multiple transformer layers (8 layers with 8 attention heads each), providing sufficient capacity to model complex regulatory dynamics while maintaining computational efficiency. The transformer processes combined representations that integrate both gene and perturbation information through a learnable fusion:

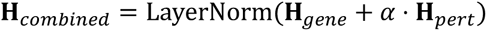

where 𝛼 is a learnable scaling parameter that balances the relative contributions of gene context and perturbation signals, allowing the model to adapt to different types of perturbations and cellular contexts.

The transformer architecture employs the following hyperparameter configuration. Each transformer layer implements a post-LayerNorm architecture where layer normalization is applied after residual connections. The feedforward network uses a 4:1 dimension expansion ratio (512 → 2048 → 512 dimensions) with GELU activation. We apply dropout regularization with a rate of 0.1 after the self-attention sublayer, within the feedforward network, and after the feedforward sublayer. The multi-head self-attention mechanism employs 8 attention heads with a head dimension of 64, and we implement memory-efficient chunked attention computation to handle large gene sets. The complete architecture consists of 8 stacked transformer layers **(Note S15**).

### 4.4 Output Prediction and Training

The final prediction layer combines the rich contextual representations learned by the transformer with the original input expression levels through a residual connection:

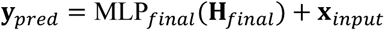

where 𝐱_𝑖𝑛𝑝𝑢𝑡_ represents the baseline gene expression. This design choice is crucial because it ensures the model learns to predict perturbation-induced changes rather than attempting to reconstruct absolute expression levels from scratch.

The residual connection serves multiple purposes: it provides a strong baseline prediction, helps with gradient flow during training, and ensures that the model’s predictions remain grounded in the observed biology of the input cells. This is particularly important when dealing with subtle perturbation effects that might only affect a small subset of genes.

For genes not present in our knowledge graph or foundation model vocabulary, we generate deterministic embeddings using gene name hashing to ensure reproducibility. While this approach allows the model to process complete gene sets, it lacks inherent biological information for truly novel genes, which represent a limitation we plan to address in future work through multi-modal information integration.

### 4.5 Training and Evaluation

Training scPert requires careful consideration of the unique challenges in perturbation response prediction. Gene expression changes are often sparse, with many genes showing little to no response to perturbations, while a smaller subset exhibits dramatic changes. Additionally, the direction of change (upregulation vs. downregulation) is often as important as the magnitude.

To address these challenges, we employ a composite loss function that combines multiple complementary objectives (**Note S14**):

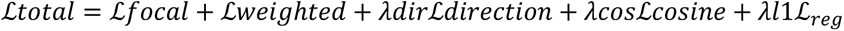

The primary component is a focal MSE loss that automatically emphasizes genes with larger expression changes:

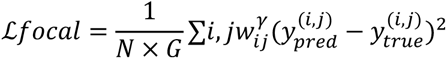

where 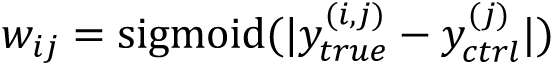 is a focal weight that increases with the magnitude of expression change, and 𝛾 is the focusing parameter. Additional loss components include direction preservation loss to maintain the sign of expression changes, cosine similarity loss for pattern consistency, and L1 regularization for model sparsity.

We evaluate model performance using complementary metrics that capture different aspects of prediction quality. Mean squared error (MSE) provides a direct measure of prediction accuracy, while Pearson correlation coefficient assesses how well the model captures the overall patterns of gene co-expression and relative changes:

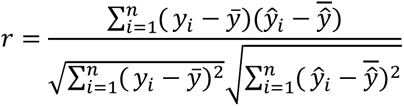

The evaluation framework includes extensive ablation studies to understand component contributions, cross-validation for generalization assessment, and comparisons with state-of-the-art baseline methods. Special attention is paid to evaluating performance on combinatorial perturbations and out-of-distribution scenarios, which represent some of the most challenging and practically relevant use cases for perturbation prediction models.

## Data Availability

For the training, testing and evaluation of scPert, all required single-cell perturbation datasets, gene embedding representations and other relevant data were sourced from publicly available databases and repositories. The following are the Gene Expression Omnibus (GEO) accession numbers and data sources used: Norman et al.: GSE133344; Adamson et al.: GSE90546; Replogle et al.: GSE146194; Dixit et al.: GSE90063; Papalexi et al.: GSE153056. Additional perturbation datasets were retrieved from https://gwps.wi.mit.edu/ (RPE-1 and K562 datasets) and https://dataverse.harvard.edu/ (processed Norman, Adamson and other datasets). The Hetionet knowledge graph used for biological knowledge integration is available at https://github.com/hetio/hetionet. scGPT pre-trained embeddings were obtained from the official scGPT repository at https://github.com/bowang-lab/scGPT. The Cancer Dependency Map (DepMap) data for cancer-relevant gene analysis were accessed from https://depmap.org/portal/. All processed datasets and gene embedding representations can be accessed through our project repository.

## Code Availability

scPert is available on GitHub (https://github.com/AI4VirtualCell/scPert) with usage documentation and comprehensive example testing datasets.

## Reference

1. Schuhmacher, A., et al. R&D efficiency of leading pharmaceutical companies - A 20-year analysis. Drug Discov Today 26, 1784–1789 (2021).

2. Pammolli, F., Magazzini, L. & Riccaboni, M. The productivity crisis in pharmaceutical R&D. Nature reviews. Drug discovery 10, 428–438 (2011).

3. Wang, K., Huang, Y., Wang, Y., You, Q. & Wang, L. Recent advances from computer-aided drug design to artificial intelligence drug design. RSC medicinal chemistry 15, 3978–4000 (2024).

4. Dixit, A. et al. Perturb-Seq: Dissecting Molecular Circuits with Scalable Single-Cell RNA Profiling of Pooled Genetic Screens. Cell 167, 1853–1866.e1817 (2016).

5. Qian, L., Dong, Z. & Guo, T. Grow AI virtual cells: three data pillars and closed-loop learning. Cell research 35, 319–321 (2025).

6. Eckert, S. et al. Decrypting the molecular basis of cellular drug phenotypes by dose-resolved expression proteomics. Nature biotechnology 43, 406–415 (2025).

7. Hwang, B., Lee, J.H. & Bang, D. Single-cell RNA sequencing technologies and bioinformatics pipelines. Experimental & molecular medicine 50, 1–14 (2018).

8. Svensson, V., Vento-Tormo, R. & Teichmann, S.A. Exponential scaling of single-cell RNA-seq in the past decade. Nat Protoc 13, 599–604 (2018).

9. Stuart, T. & Satija, R. Integrative single-cell analysis. Nat Rev Genet 20, 257–272 (2019).

10. Yao, D. et al. Scalable genetic screening for regulatory circuits using compressed Perturb-seq. Nature biotechnology 42, 1282–1295 (2024).

11. Wessels, H.-H. et al. Efficient combinatorial targeting of RNA transcripts in single cells with Cas13 RNA Perturb-seq. Nature methods 20, 86–94 (2023).

12. Liu, N. et al. Scalable, compressed phenotypic screening using pooled perturbations. Nature biotechnology 43, 1324–1336 (2024).

13. Moeckel, C., Mouratidis, I., Chantzi, N., Uzun, Y. & Georgakopoulos-Soares, I. Advances in computational and experimental approaches for deciphering transcriptional regulatory networks: Understanding the roles of cis-regulatory elements is essential, and recent research utilizing MPRAs, STARR-seq, CRISPR-Cas9, and machine learning has yielded valuable insights. BioEssays: news and reviews in molecular, cellular and developmental biology 46, e2300210 (2024).

14. Roohani, Y., Huang, K. & Leskovec, J. Predicting transcriptional outcomes of novel multigene perturbations with GEARS. Nature biotechnology 42, 927–935 (2024).

15. Lotfollahi, M. et al. Predicting cellular responses to complex perturbations in high-throughput screens. Mol Syst Biol 19, e11517 (2023).

16. Replogle, J.M. et al. Mapping information-rich genotype-phenotype landscapes with genome-scale Perturb-seq. Cell 185, 2559–2575.e2528 (2022).

17. Alsalloum, G.A., Al Sawaftah, N.M., Percival, K.M. & Husseini, G.A. Digital Twins of Biological Systems: A Narrative Review. IEEE open journal of engineering in medicine and biology 5, 670–677 (2024).

18. De Domenico, M. et al. Challenges and opportunities for digital twins in precision medicine from a complex systems perspective. NPJ digital medicine 8, 37 (2025).

19. Cui, H. et al. scGPT: toward building a foundation model for single-cell multi-omics using generative AI. Nat Methods 21, 1470–1480 (2024).

20. Hao, M. et al. Large-scale foundation model on single-cell transcriptomics. Nat Methods 21, 1481–1491 (2024).

21. Yang, X. et al. GeneCompass: deciphering universal gene regulatory mechanisms with a knowledge-informed cross-species foundation model. Cell Res 34, 830–845 (2024).

22. Zeng, Y. et al. CellFM: a large-scale foundation model pre-trained on transcriptomics of 100 million human cells. Nat Commun 16, 4679 (2025).

23. Istrate, A.-M., Li, D. & Karaletsos, T. scGenePT: Is language all you need for modeling single-cell perturbations? bioRxiv (2024). 10.1101/2024.10.23.619972

24. Chen, Y. & Zou, J. GenePert: Leveraging GenePT Embeddings for Gene Perturbation Prediction. bioRxiv (2024). 10.1101/2024.10.27.620513

25. Zhu, O. & Li, J. Scouter predicts transcriptional responses to genetic perturbations with large language model embeddings. Nat Comput Sci (2025). 10.1038/s43588-025-00912-8

26. Chen, Y. & Zou, J. Simple and effective embedding model for single-cell biology built from ChatGPT. Nat Biomed Eng 9, 483–493 (2025).

27. He, S. et al. Squidiff: predicting cellular development and responses to perturbations using a diffusion model. Nat Methods 23, 65–77 (2026).

28. Ben Guebila, M., et al. GRAND: a database of gene regulatory network models across human conditions. Nucleic acids research 50, D610–D621 (2022).

29. Szklarczyk, D. et al. The STRING database in 2023: protein-protein association networks and functional enrichment analyses for any sequenced genome of interest. Nucleic acids research 51, D638–D646 (2023).

30. Kanehisa, M., Furumichi, M., Sato, Y., Kawashima, M. & Ishiguro-Watanabe, M. KEGG for taxonomy-based analysis of pathways and genomes. Nucleic Acids Res 51, D587–D592 (2023).

31. Mao, G. et al. Predicting gene regulatory links from single-cell RNA-seq data using graph neural networks. Briefings in bioinformatics 24, bbad414 (2023).

32. Tan, Y. et al. BioDSNN: a dual-stream neural network with hybrid biological knowledge integration for multi-gene perturbation response prediction. Brief Bioinform 26, bbae617 (2024).

33. Piran, Z., Cohen, N., Hoshen, Y. & Nitzan, M. Disentanglement of single-cell data with biolord. Nat Biotechnol 42, 1678–1683 (2024).

34. Bunne, C. et al. Learning single-cell perturbation responses using neural optimal transport. Nat Methods 20, 1759–1768 (2023).

35. Wei, Z. et al. Benchmarking algorithms for generalizable single-cell perturbation response prediction. Nat Methods (2025). 10.1038/s41592-025-02980-0

36. Ahlmann-Eltze, C., Huber, W. & Anders, S. Deep-learning-based gene perturbation effect prediction does not yet outperform simple linear baselines. Nat Methods 22, 1657–1661 (2025).

37. Zhou, Y. et al. TTD: Therapeutic Target Database describing target druggability information. Nucleic acids research 52, D1465–D1477 (2024).

38. Himmelstein, D.S. et al. Systematic integration of biomedical knowledge prioritizes drugs for repurposing. eLife 6, e26726 (2017).

39. Theodoris, C.V. et al. Transfer learning enables predictions in network biology. Nature 618, 616–624 (2023).

40. Csendes, G., Sanz, G., Szalay, K.Z. & Szalai, B. Benchmarking foundation cell models for post-perturbation RNA-seq prediction. BMC genomics 26, 393 (2025).

41. Norman, T.M. et al. Exploring genetic interaction manifolds constructed from rich single-cell phenotypes. Science 365, 786–793 (2019).

42. Adamson, B. et al. A Multiplexed Single-Cell CRISPR Screening Platform Enables Systematic Dissection of the Unfolded Protein Response. Cell 167, 1867–1882.e1821 (2016).

43. Vinas Torne, R., et al. Systema: a framework for evaluating genetic perturbation response prediction beyond systematic variation. Nat Biotechnol (2025). 10.1038/s41587-025-02777-8

44. Aibar, S. et al. SCENIC: single-cell regulatory network inference and clustering. Nature methods 14, 1083–1086 (2017).

45. Liu, Y., Su, Z., Tavana, O. & Gu, W. Understanding the complexity of p53 in a new era of tumor suppression. Cancer cell 42, 946–967 (2024).

46. Hopkins, A.L. Network pharmacology: the next paradigm in drug discovery. Nature chemical biology 4, 682–690 (2008).

47. AbdulAzeez, S. & Borgio, J.F. In-Silico Computing of the Most Deleterious nsSNPs in HBA1 Gene. PloS one 11, e0147702 (2016).

48. Ma, W.K. et al. ASO-Based PKM Splice-Switching Therapy Inhibits Hepatocellular Carcinoma Growth. Cancer research 82, 900–915 (2022).

49. Shih, Y.-T. & Hsueh, Y.-P. The involvement of endoplasmic reticulum formation and protein synthesis efficiency in VCP- and ATL1-related neurological disorders. Journal of biomedical science 25, 2 (2018).

50. Baumgartner, C.K. et al. The PTPN2/PTPN1 inhibitor ABBV-CLS-484 unleashes potent anti-tumour immunity. Nature 622, 850–862 (2023).

51. Stracquadanio, G. et al. The importance of p53 pathway genetics in inherited and somatic cancer genomes. Nature reviews. Cancer 16, 251–265 (2016).

52. Milacic, M. et al. The Reactome Pathway Knowledgebase 2024. Nucleic acids research 52, D672–D678 (2024).

53. Pabst, T. et al. Dominant-negative mutations of CEBPA, encoding CCAAT/enhancer binding protein-alpha (C/EBPalpha), in acute myeloid leukemia. Nature genetics 27, 263–270 (2001).

54. Ji, Z. et al. The forkhead transcription factor FOXK2 promotes AP-1-mediated transcriptional regulation. Molecular and cellular biology 32, 385–398 (2012).

55. Allouch, A. et al. CDKN1A is a target for phagocytosis-mediated cellular immunotherapy in acute leukemia. Nature communications 13, 6739 (2022).

56. Dadsena, S. et al. Lipid unsaturation promotes BAX and BAK pore activity during apoptosis. Nature communications 15, 4700 (2024).

57. Kamimura, R. et al. Identification of Binding Proteins for TSC22D1 Family Proteins Using Mass Spectrometry. International journal of molecular sciences 22 (2021).

58. Zhang, J., Dang, F., Ren, J. & Wei, W. Biochemical Aspects of PD-L1 Regulation in Cancer Immunotherapy. Trends in biochemical sciences 43, 1014–1032 (2018).

59. Papalexi, E. et al. Characterizing the molecular regulation of inhibitory immune checkpoints with multimodal single-cell screens. Nature genetics 53, 322–331 (2021).

60. Mimitou, E.P. et al. Multiplexed detection of proteins, transcriptomes, clonotypes and CRISPR perturbations in single cells. Nature methods 16, 409–412 (2019).

61. Tsherniak, A. et al. Defining a Cancer Dependency Map. Cell 170, 564–576.e516 (2017).

62. Pacini, C. et al. A comprehensive clinically informed map of dependencies in cancer cells and framework for target prioritization. Cancer cell 42, 301–316.e309 (2024).

63. Sun, X. et al. PROTACs: great opportunities for academia and industry. Signal transduction and targeted therapy 4, 64 (2019).

64. Pundhir, S. et al. Enhancer and Transcription Factor Dynamics during Myeloid Differentiation Reveal an Early Differentiation Block in Cebpa null Progenitors. Cell reports 23, 2744–2757 (2018).

65. Verbiest, T., Bouffler, S., Nutt, S.L. & Badie, C. PU.1 downregulation in murine radiation-induced acute myeloid leukaemia (AML): from molecular mechanism to human AML. Carcinogenesis 36, 413–419 (2015).

66. Li, Q. et al. PRDM1/BLIMP1 induces cancer immune evasion by modulating the USP22-SPI1-PD-L1 axis in hepatocellular carcinoma cells. Nature communications 13, 7677 (2022).

67. Stankey, C.T. et al. A disease-associated gene desert directs macrophage inflammation through ETS2. Nature 630, 447–456 (2024).

68. Wang, G. et al. Identification of endothelial and mesenchymal FOXF1 enhancers involved in alveolar capillary dysplasia. Nature communications 15, 5233 (2024).

69. Migale, R. et al. FOXL2 interaction with different binding partners regulates the dynamics of ovarian development. Science advances 10, eadl0788 (2024).

70. Schmidt, V., Sieckmann, T., Kirschner, K.M. & Scholz, H. WT1 regulates HOXB9 gene expression in a bidirectional way. Biochimica et biophysica acta. Gene regulatory mechanisms 1864, 194764 (2021).

71. Douglas, N.C. & Papaioannou, V.E. The T-box transcription factors TBX2 and TBX3 in mammary gland development and breast cancer. Journal of mammary gland biology and neoplasia 18, 143–147 (2013).

72. Romanova, E.I. et al. RUNX1/CEBPA mutation in acute myeloid leukemia promotes hypermethylation and indicates for demethylation therapy. International journal of molecular sciences 23, 11413 (2022).

73. Grossmann, V. et al. Expression of CEBPA is reduced in RUNX1-mutated acute myeloid leukemia. Blood cancer journal 2, e86 (2012).

74. Wu, J. et al. Biology-driven insights into the power of single-cell foundation models. Genome Biol 26, 334 (2025).

75. Zhang, M. et al. GREmLN: a cellular regulatory network-aware transcriptomics foundation model. bioRxiv (2025). 10.1101/2025.07.03.663009

76. Fang, G. et al. Discovering genetic interactions bridging pathways in genome-wide association studies. Nat Commun 10, 4274 (2019).

77. Costanzo M. et al. A global genetic interaction network maps a wiring diagram of cellular function. Science 353, aaf1420 (2016).

78. Ngoi NYL. et al. Synthetic lethal strategies for the development of cancer therapeutics. Nat Rev Clin Oncol. 22, 46–64 (2025).

79. Lian, X. et al. SingPro: a knowledge base providing single-cell proteomic data. Nucleic Acids Res 52, D552–D561 (2024).

80. UniProt Consortium. UniProt: the Universal Protein Knowledgebase in 2023. Nucleic Acids Res. 51, D523–D531 (2023).

81. Szklarczyk D. et al. The STRING database in 2023: protein–protein association networks and functional enrichment analysis for any sequenced genome of interest. Nucleic Acids Res. 51, D638–D646 (2023).

82. Vashishth, S., Sanyal, S., Nitin, V. & Talukdar, P. Composition-based Multi-Relational Graph Convolutional Networks. in International Conference on Learning Representations (2020).

