## Supplemental File 1 for "A Multi-modal LLM-Knowledge Fusion Framework for Predicting Single-cell Genetic Perturbation Effects"

### Supplementary Notes

Note S1: Dataset characteristics and preprocessing procedures

Note S2: Comparative analysis of single-cell perturbation response prediction methods

Note S3: An introduction of the Pearson correlation delta used in this study

Note S4: An introduction of the Mean Squared Error used in this study

Note S5: Centroid accuracy for evaluating perturbation prediction

Note S6: Genetic interaction classifications

Note S7: Precision@10 metric for genetic interaction evaluation

Note S8: Genetic interaction scoring systems and classification thresholds

Note S9: Model evaluation for predicting genetic interactions

Note S10: Performance evaluation on p53 pathway perturbations

Note S11: Mixscape-based quality control and perturbation validation

Note S12: Prediction quality score calculation methodology

Note S13: CompGCN integration and ablation analysis

Note S14: Loss function of scPert

Note S15: Hyperparameters of scPert

### Supplementary Figures

Figure S1: Statistical overview of perturbation datasets used for model evaluation

Figure S2: Detailed statistical characterization of benchmark datasets

Figure S3: Prediction of scPert for Top 20 DE genes in different perturbations

Figure S4: Comprehensive evaluation of scPert prediction quality across perturbation conditions and target genes

Figure S5: CEBPA-dominant perturbation effects on gene expression

Figure S6: UMAP representation and distance correlation analysis of single versus combination perturbations

### Supplementary Tables

Table S1: Statistical overview of perturbation datasets used for model evaluation

Table S2: Mean squared error performance of single gene perturbation

Table S3: Pearson correlation performance of single gene perturbation

Table S4: Centroid accuracy performance of single gene perturbation

Table S5: Mean squared error performance of combinatorial perturbation

Table S6: Pearson correlation performance of combinatorial perturbation

Table S7: Precision@10 scores for genetic interaction prediction

Table S8: Performance evaluation on p53-associate gene

Table S9: Mean squared error performance of ablation experiment

Table S10. Ablation experiments on centroid accuracy

### Supplementary Notes

#### Note S1: Dataset characteristics and preprocessing procedures

The comprehensive evaluation of perturbation prediction methods requires diverse datasets spanning different cell types, experimental scales, and perturbation modalities. Our analysis incorporated five major single-cell perturbation datasets that collectively represent the current landscape of high-throughput genetic screens. The Norman[1] dataset contains gene expression data from 131 two-gene perturbations and 105 one-gene perturbations, with each perturbation condition replicated across approximately 300-700 cells. This dataset serves as a benchmark for combinatorial perturbation prediction due to its systematic coverage of pairwise gene interactions and robust experimental design. The Adamson[2] dataset, also utilizing K562 cells with Perturb-seq, focuses on single-gene perturbations with 87 unique CRISPR interference targets, each replicated in around 100 cells, providing a complementary perspective on individual gene function within the same cellular context.

The Replogle[3] dataset represents the largest scale perturbation screen in our analysis, encompassing genome-wide CRISPR interference perturbations in both K562 leukemia and RPE-1 epithelial cell lines. Following quality control procedures that retained perturbations with strong transcriptional phenotypes, the processed dataset consists of 162,751 samples from 1,092 perturbations. This scale enables robust statistical analysis while providing sufficient coverage of diverse gene functions. The Dixit[4] dataset, accessed through GEO accession GSE90063, provides additional validation data for single-gene perturbations, while the supplementary RPE-1 dataset from the Replogle collection offers cross-cell-type validation of perturbation effects.

Data preprocessing followed standardized protocols to ensure comparability across datasets and experimental platforms. First, gene identifier harmonization was performed to address nomenclature inconsistencies across sources: dataset-specific identifiers (e.g., Ensembl IDs, outdated symbols) were mapped to official HGNC symbols, then aligned to the scGPT model's canonical vocabulary. For perturbation conditions, biological equivalence was standardized: combinatorial perturbations (e.g., GeneA+GeneB vs. GeneB+GeneA) were sorted alphabetically to unify identifiers, and all control types (e.g., non-targeting guides, safe-harbor perturbations) were consolidated into a single "control" designation after verifying consistent expression profiles. Gene expression values were normalized per cell using total count normalization followed by log transformation to stabilize variance and approximate normal distributions.

Highly variable genes were selected using established criteria, typically retaining 2,000-5,000 genes per dataset depending on the experimental scale and sequencing depth. Perturbation-specific genes were always included in the analysis regardless of their variance characteristics to maintain biological relevance. Quality control metrics included removal of cells with extremely low or high gene counts, mitochondrial gene percentage filtering, and doublet detection using computational methods. For combinatorial perturbation experiments, additional validation steps confirmed successful perturbation of all intended targets through direct measurement of target gene expression levels. All processed datasets can be downloaded using GEARS API.

#### Note S2: Comparative analysis of single-cell perturbation response prediction methods

Recent advances in perturbation prediction have explored diverse strategies for incorporating biological knowledge and leveraging large-scale models. We compare scPert with five representative approaches that employ distinct methodological paradigms: GEARS[5], a knowledge graph-enhanced approach that represents the current state-of-the-art in perturbation prediction; scGPT[6], a foundation model specifically designed for single-cell genomics; Biolord[7], a deep generative framework that disentangles single-cell multi-omic representations into known and unknown biological attributes and uses this disentangled latent space to generate counterfactual cellular profiles; scGenePT[8], which augments scGPT with additional gene language representations derived from unstructured biological knowledge; and Scouter[9], a method that uses large language model-derived gene embeddings combined with a compressor–generator neural network to predict transcriptional responses.

The GEARS framework introduces distinct gene and perturbation embeddings that capture different aspects of cellular response. Gene embeddings encode intrinsic properties such as baseline expression patterns and regulatory roles, while perturbation embeddings represent the specific effects of disrupting gene function. This separation enables the model to distinguish between a gene's normal cellular role and its perturbation phenotype, a critical distinction for accurate prediction. The compositional operator that combines these embeddings with cross-gene information allows GEARS to predict complex combinatorial effects by modeling how multiple perturbations interact within the cellular network. Experimental validation has demonstrated GEARS's superior performance in predicting genetic interaction subtypes including synergy, suppression, neomorphism, redundancy, and epistasis, achieving correlation coefficients of approximately 0.4 for most interaction types compared to near-zero performance from naive approaches.

scGPT represents the foundation model approach to single-cell genomics, leveraging transformer architectures pretrained on massive datasets to capture general patterns in gene expression. The model processes single-cell data by treating genes as tokens and cells as sequences, enabling it to learn complex dependencies between genes through self-attention mechanisms. This approach excels at capturing global expression patterns and has demonstrated strong performance across diverse tasks including cell type annotation, batch integration, and reference mapping. However, the foundation model paradigm faces inherent challenges in perturbation prediction tasks due to the domain shift between natural gene expression patterns and perturbation-induced changes.

Biolord employs a deep generative framework that explicitly disentangles single-cell data into known attributes (cell type, perturbation, temporal state) and unknown attributes through variational autoencoders. The model implements latent optimization, randomly initializing latent codes rather than using encoder networks, which prevents the entanglement issues that plague amortized inference approaches. The architecture constructs a decomposed latent space with dedicated subnetworks for each attribute: categorical attributes use shared embeddings optimized directly, while ordered attributes employ multilayer perceptrons to capture structural information. However, biolord does not explicitly incorporate structured biological knowledge such as gene regulatory networks or pathway information, instead relying on training data to implicitly learn these relationships through the joint optimization of decomposed representations and generative objectives.

scGenePT extends the scGPT foundation model by augmenting gene representations with language model embeddings derived from NCBI gene descriptions and Gene Ontology annotations, bridging data-driven learning with textual biological knowledge. The model maintains scGPT's transformer encoder-decoder architecture but adds a dedicated embedding layer for language-derived features, with each gene receiving four representations: expression counts, gene tokens, perturbation tokens, and language embeddings generated using OpenAI's text-embedding models. These separate embeddings are element-wise summed to create unified gene representations fed into the transformer architecture. During fine-tuning for perturbation prediction, gene token and expression count embeddings initialize from pretrained scGPT weights while language and perturbation embeddings train from scratch, creating a supervised alignment process. While this successfully incorporates textual biological knowledge, the additive fusion strategy treats all modalities equally without learning optimal weighting schemes. The simple element-wise summation does not allow dynamic adjustment of relative importance between structured knowledge and data-driven representations based on prediction context, potentially limiting adaptability across diverse perturbation scenarios where the complementarity between expression-based and text-based information varies substantially.

Scouter represents a distinctly lightweight approach leveraging gene embeddings from ChatGPT model applied to NCBI gene descriptions, implementing a compressor-generator framework where control cell expression is reduced to compact latent representations, concatenated with fixed LLM embeddings, and fed to a generator producing perturbation predictions. Scouter's

1,536-dimensional LLM embeddings remain completely fixed during training, serving as pretrained priors encoding gene functions and regulatory relationships extracted from textual descriptions. This eliminates embedding alignment challenges but sacrifices adaptability to dataset-specific patterns or cell-type-specific behaviors. However, fixed embeddings prevent adaptation to nuanced dataset-specific regulatory patterns, and while simple concatenation overcomes GO graph coverage limitations, it lacks explicit gene-gene interaction modeling through structured knowledge graphs or rich contextualization from massive pretraining that characterizes foundation model approaches.

scPert's architectural innovations position it as a complementary advance that bridges multiple methodological paradigms. Unlike biolord's purely generative approach, scPert directly integrates curated biological knowledge through knowledge graph embeddings processed via CompGCN, providing explicit encoding of gene-disease associations, pathway memberships, and molecular function relationships. This structured knowledge integration complements rather than replaces data-driven representations from foundation models, addressing a key limitation of methods that rely solely on implicit learning. Compared to scGenePT's additive fusion and Scouter's fixed embeddings, scPert's hierarchical fusion mechanism learns optimal combination strategies during training, allowing the model to adaptively weight different modalities. The memory-efficient transformer architecture further distinguishes scPert by explicitly modeling gene-gene interactions through multi-head attention mechanisms that adapt to specific biological contexts, rather than relying solely on compressed latent representations or fixed pretrained embeddings. This design philosophy aligns with emerging insights suggesting that hybrid approaches combining multiple complementary information sources including structured knowledge graphs, foundation model representations, and biological priors, can achieve superior generalization compared to methods relying on single modalities or simple combination strategies.

#### Note S3. An introduction of the *Pearson correlation delta* used in this study

Accurate evaluation of perturbation prediction methods requires metrics that specifically capture the biological phenomena of interest rather than general expression similarity. The Pearson correlation delta (Pearson_delta) metric addresses fundamental limitations of conventional evaluation approaches by focusing on perturbation-induced changes rather than absolute expression values. This metric calculates the correlation between predicted and observed expression changes relative to control conditions, effectively measuring the model's ability to capture the perturbation response signature while controlling for baseline expression differences across cells and experimental conditions.

Conventional Pearson correlation metrics can be confounded by high baseline expression similarity between perturbed and control conditions, potentially yielding misleadingly high correlations even when perturbation-specific effects are poorly predicted. By computing correlations on the delta (Δ) values representing the difference between perturbed and control expression profiles, Pearson_delta isolates the perturbation-induced transcriptional response, ensuring that model performance reflects genuine predictive capability for intervention effects rather than merely recapitulating unperturbed expression patterns.

Mathematically, the Pearson correlation delta (Pearson_delta) is defined as follows:

$$Pearson\_delta = \frac{\mathrm{cov}(\Delta\mathbf{y}_{pred}, \Delta\mathbf{y}_{true})}{\sigma_{\Delta\mathbf{y}_{pred}}\sigma_{\Delta\mathbf{y}_{true}}}$$

Where $\Delta\mathbf{y} = \mathbf{y}_{perturbed} - \bar{\mathbf{y}}_{control}$ represents the gene expression change relative to the mean of the control group. This formulation ensures that the metric captures the perturbation-induced transcriptional response signature rather than absolute expression levels.

Furthermore, the delta-based correlation framework naturally accounts for technical and biological variability inherent in single-cell data, including batch effects and cell-type-specific baseline expression heterogeneity. By centering evaluation on relative changes, Pearson_delta provides a more biologically interpretable assessment of whether models capture the causal mechanisms underlying perturbation responses, the directional shifts in gene regulatory networks that define cellular phenotypic transitions. This metric alignment with mechanistic understanding makes it superior for evaluating models intended to predict intervention outcomes in drug discovery, disease modeling, and precision medicine applications where perturbation effect prediction, not steady-state reconstruction, is the primary objective.

#### Note S4. An introduction of the *Mean Squared Error* used in this study

Mean squared error (MSE) can be naturally extended to the delta expression space to provide a robust quantitative assessment of perturbation prediction accuracy. In this setting, MSE is computed between predicted and observed expression changes relative to the corresponding control condition, rather than on absolute expression levels. By operating in the delta space, this metric explicitly focuses on perturbation-induced effects while controlling for baseline expression variability across cells, genes, and experimental conditions. The squared error formulation penalizes large discrepancies in the magnitude of predicted responses, making delta-space MSE particularly sensitive to failures in capturing the strength of gene-level regulation induced by perturbations. As a result, delta-space MSE offers a complementary perspective to correlation-based metrics by directly measuring absolute deviations in perturbation response intensity.

To provide a robust quantitative assessment of absolute deviations in the delta expression space, MSE is calculated as:

$$MSE_{\Delta} = \frac{1}{G}\sum_{j=1}^{G} (\Delta y_{pred,j} - \Delta y_{true,j})^2$$

Where $G$ denotes the number of genes analyzed, and $\Delta y_j$ represents the expression change of gene $j$ relative to the control. This metric is particularly sensitive to discrepancies in the magnitude of predicted responses, complementing correlation-based evaluations.

#### Note S5: Centroid accuracy for evaluating perturbation prediction

Centroid accuracy is used to evaluate whether a model correctly captures the perturbation-specific expression signature at the population level. For a given perturbation $X$, we first compute the predicted centroid $O_{\text{pred}}(X)$and the corresponding ground-truth centroid $O(X)$, each representing the average expression profile of cells under perturbation $X$in the embedding or gene expression space. Centroid accuracy measures the proportion of other perturbations $Y \in \mathcal{P}$for which the predicted centroid of $X$is closer to its own ground-truth centroid than to the ground-truth centroids of alternative perturbations. Formally, this is quantified using an indicator function that compares Euclidean distances between centroids. A higher centroid accuracy indicates that the model not only predicts expression changes with small error, but also preserves perturbation identity by placing predicted responses closer to their true perturbation-specific manifold than to unrelated perturbations. This metric therefore captures the discriminability of perturbation effects in the learned representation space, complementing error- and correlation-based evaluations.

$$\text{Centroid accuracy}(X) = \frac{1}{|\mathcal{P}| - 1} \sum_{Y \in \mathcal{P}} \mathbb{1}\left[d\left(O_{\text{pred}}(X), O(X)\right) < d\left(O_{\text{pred}}(X), O(Y)\right)\right]$$

#### Note S6: Genetic interaction classifications

To identify and categorize genetic interaction subtypes, we follow the classification framework and scoring metrics defined by Norman et al. and later adopted by Roohani et al. in the GEARS model. This established framework provides standardized criteria for distinguishing different interaction modalities through quantitative analysis of combinatorial perturbation effects relative to individual gene perturbations. By using this existing classification scheme instead of developing new criteria, we ensure direct comparability of our results with prior studies and enable meaningful benchmarking against established methods. The framework classifies genetic interactions into five principal categories based on their deviation from simple additive effects: originally, Norman et al. defined seven types (additive, epistatic, neomorphic, potentiation, redundant, suppressive, and synergistic), but Roohani et al. streamlined this taxonomy by merging synergistic and potentiation categories, as both represent cases where combinatorial effects exceed the sum of individual perturbations, with differences that are quantitative rather than qualitatively distinct in mechanism. We adopt GEARS's five-category system (synergy, suppression, neomorphism, redundancy, and epistasis) as it balances biological interpretability with reliable classification in practice.

Synergistic interactions occur when the combined effect of perturbing two genes exceeds the sum of their individual effects, quantified by magnitude scores above 1.15 in the GEARS scoring system. These interactions often involve genes acting in parallel pathways that converge on common cellular processes; when both pathways are disrupted simultaneously, compensatory mechanisms that buffer single-pathway perturbations become insufficient, leading to amplified cellular responses beyond simple additive predictions. This synergy reflects the cell's inability to maintain homeostasis when multiple redundant or parallel control systems are compromised at the same time, revealing functional relationships where pathway-level redundancy provides robustness against individual perturbations but fails under combinatorial disruption. Suppression interactions, by contrast, happen when one gene's perturbation masks or reduces the effect of another, identified by magnitude scores below 1.0 (indicating the combinatorial effect is weaker than expected from summing individual effects). These typically occur in hierarchically organized pathways where downstream components functionally constrain upstream regulatory inputs, revealing pathway structure and rate-limiting steps where cellular responses are bottlenecked by specific molecular functions regardless of upstream activation status. Understanding suppression patterns helps identify dominant pathway branches and critical control points in regulatory networks, distinguishing genes whose perturbation effects can be compensated from those representing non-redundant pathway components.

Neomorphic interactions produce entirely novel phenotypes not seen with either individual perturbation, identified computationally by low model fit scores below 0.88, which signal that linear additive models cannot explain the combinatorial response. These interactions represent cellular states outside the normal physiological range, often arising from disrupted balanced regulatory networks that maintain homeostasis. Neomorphic phenotypes emerge when opposing regulatory forces are removed simultaneously, creating novel expression patterns that would not occur through modifying individual pathway components; they offer opportunities for therapeutic intervention via combination approaches that exploit emergent vulnerabilities absent under single-perturbation conditions, particularly relevant for synthetic lethality strategies in cancer treatment. Redundancy interactions occur when genes perform overlapping functions, so individual perturbations have minimal effects while combined perturbation reveals the full phenotypic consequence, which is quantified by similarity scores above 0.85 between individual and combinatorial effects. This category is key for understanding genetic robustness mechanisms and identifying potential therapeutic targets where single-agent approaches may fail due to functional compensation: redundant gene pairs act as evolutionarily conserved backup systems protecting cells from loss of critical functions, indicating cells have invested evolutionary resources in maintaining alternative pathways for essential processes, processes under strong selective pressure that may become vulnerable dependencies when all redundant pathways are targeted together.

Epistatic interactions involve one gene masking the phenotypic expression of another, typically when genes function in the same linear pathway where the downstream gene's function is required to observe the upstream gene's effect. These are identified by equality of contribution scores above 0.28, reflecting asymmetric contributions where one gene dominates the combinatorial response. Epistatic relationships define pathway architecture by identifying which genes act upstream versus downstream in regulatory cascades, and understanding them is essential for predicting combinatorial perturbation outcomes and identifying pathway bottlenecks where therapeutic intervention is most effective, since targeting downstream epistatic suppressors may bypass upstream regulatory dysfunction.

We use the classification thresholds established by Roohani et al. in the GEARS framework without modification: these values have been empirically validated across multiple perturbation datasets and serve as robust criteria for distinguishing interaction modalities in single-cell perturbation experiments. Balancing sensitivity and specificity, the thresholds ensure classifications capture genuine biological phenomena while minimizing false positives from technical noise or insufficient statistical power. Specifically, we apply magnitude thresholds of >1.15 for synergy and .0 for suppression, a model fit threshold of .88 for neomorphism, a similarity threshold of >0.85 for redundancy, and an equality of contribution threshold of >0.28 for epistasis. These thresholds were selected by GEARS developers through systematic analysis of experimentally validated genetic interactions, maximizing agreement between computational predictions and experimental observations. By retaining identical threshold criteria, our evaluation ensures that differences in genetic interaction prediction performance between scPert and baseline methods reflect true differences in model capability rather than artifacts from inconsistent classification schemes. While alternative thresholds have been proposed in the literature (with some studies using stricter or more permissive criteria based on experimental context and false positive tolerance), adopting GEARS's thresholds facilitates direct comparison with the current state-of-the-art method and enables meta-analyses combining results across studies using this standardized framework. Future work may explore adaptive threshold selection that accounts for dataset-specific characteristics like perturbation efficiency, cellular heterogeneity, and measurement noise, but our current analysis prioritizes consistency with established benchmarks to enable rigorous comparative evaluation.

#### Note S7: Precision@10 metric for genetic interaction evaluation

Precision@10 is a critical evaluation metric for assessing the practical value of genetic interaction prediction methods, especially in experimental settings where validation resources are limited. It directly addresses the real-world need for researchers to prioritize a small subset of predicted interactions for follow-up experiments, making the accuracy of top-ranked predictions pivotal to successful biological discovery. Specifically, this metric quantifies the proportion of the top 10 predicted genetic interactions that are experimentally validated as true biological interactions, offering a stringent measure of model performance that directly translates to experimental success rates.

The rationale for Precision@10 stems from the combinatorial complexity of genetic interactions: the number of potential gene pairs grows quadratically with the number of genes under study. For example, a screen of 100 genes yields nearly 5,000 pairwise combinations, rendering exhaustive validation infeasible. Since researchers can only validate a small fraction of predictions, metrics focusing on top-ranked quality instead of overall classification accuracy are essential. Precision@10 fills this gap by estimating the expected success rate of validating a model's highest-confidence predictions, providing actionable guidance for experimental prioritization.

Computationally, Precision@10 works by ranking all possible gene pairs based on their predicted scores for a specific interaction subtype, selecting the top 10 highest-scoring candidates, and calculating the fraction of these that align with experimentally confirmed interactions (as defined by the classification framework in Note S6). Mathematically, for a given genetic interaction subtype $s$, $\text{Precision@10}_s$ is defined as = (number of true interactions in top 10 predictions) / 10. Following the GEARS implementation, we compute this metric separately for each of the five genetic interaction subtypes (synergy, suppression, neomorphism, redundancy, epistasis) instead of aggregating results. This subtype-specific approach is necessary because different interaction types have distinct baseline frequencies in datasets and rely on unique biological signatures. For instance, synergy involves amplified magnitude, epistasis involves asymmetric gene contributions, and neomorphism involves deviation from additive models. Separating scores enables granular assessment of whether models capture each interaction modality effectively or exhibit biases toward specific types.

Interpretation of Precision@10 is straightforward: values near 1.0 indicate nearly all top predictions are true interactions, supporting immediate validation; values around 0.5 suggest moderate reliability, warranting additional computational filtering; and values substantially below 0.5 signal insufficient prediction quality for experimental guidance without refinement. The metric has inherent limitations, however: for rare interaction subtypes with fewer than 10 true positives in the dataset, the maximum achievable Precision@10 is constrained by the number of available valid interactions, requiring context-dependent interpretation. Additionally, while the threshold of 10 predictions is practically motivated by typical validation scales, it is an arbitrary cutoff that can be adjusted based on a laboratory's specific capabilities and resources.

#### Note S8: Genetic interaction scoring systems and classification threshold

The quantitative assessment of genetic interactions employs a mathematical framework based on linear models to identify deviations from additive perturbation effects[5]. For a cell with k genes under perturbation, the post-perturbation gene expression vector $\delta^{ab}$ represents the magnitude of mean gene expression changes compared to unperturbed control cells, calculated as $\delta^{ab} = \bar{x}^{ab} - \bar{x}^{c}$, where $\bar{x}^{ab}$ denotes the mean gene expression value of perturbed cells and $\bar{x}^{c}$ represents the corresponding control expression levels. The mathematical foundation for genetic interaction detection relies on fitting linear models to predict combinatorial effects based on individual perturbation responses: $\delta^{ab} = \alpha + \beta\cdot\delta^{a} + \gamma\cdot\delta^{b} + \varepsilon$, where the error term $\varepsilon$ captures deviations from additivity that indicate genetic interactions.

| GI Score | Mathematical Definition | Biological Interpretation |
|---|---|---|
| Model fit | $\mathrm{corr}(\alpha + \beta\cdot\delta^{a} + \gamma\cdot\delta^{b}, \delta^{ab})$ | Measures how well additive model explains combinatorial effects |
| Magnitude | $\sqrt{(\alpha^2 + \beta^2)}$ | Quantifies overall interaction strength |
| Similarity | $\mathrm{corr}([\delta^{a}, \delta^{b}], \delta^{ab})$ | Assesses resemblance between single and combinatorial effects |
| Equality of contribution | $\min(\mathrm{corr}(\delta^{a},\delta^{ab}),\mathrm{corr}(\delta^{b},\delta^{ab}))$ $/\max(\mathrm{corr}(\delta^{a},\delta^{ab}),\mathrm{corr}(\delta^{b},\delta^{ab}))$ | Determines balance of individual gene contributions |

These four complementary scores capture different aspects of genetic interactions and are computed independently for each combinatorial perturbation. The model fit score assesses linearity of the combinatorial response, with low values indicating that simple additive models fail to explain observed effects. The magnitude score quantifies interaction strength in absolute terms, with values above 1.0 suggesting amplification and values below 1.0 suggesting dampening relative to individual effects. The similarity score compares the combinatorial expression profile to individual perturbation profiles through concatenation of single-gene delta vectors, identifying cases where combinations resemble one or both individual perturbations. The equality of contribution score employs a min-max ratio to detect asymmetry, with values near 0 indicating complete dominance by one gene and values near 1 indicating balanced contributions.

| Interaction Type | Threshold Criterion | Definition |
|---|---|---|
| Synergy | Magnitude > 1.15 | Combined effect exceeds sum of individual effects |
| Suppressive | Magnitude < 1.0 | One perturbation dampens the effect of another |
| Neomorphism | Model fit < 0.88 | Novel phenotypes emerge from gene combination |
| Redundant | Similarity > 0.85 | Functional overlap between targeted genes |
| Epistasis | Equality of contribution > 0.28 | Asymmetric contributions with one gene dominating |

The classification thresholds employed in this study follow the established criteria from the GEARS framework to ensure consistency with previous genetic interaction analyses and enable direct comparison of results across different prediction methods. These threshold values have been empirically validated across multiple datasets and represent robust criteria for distinguishing between different genetic interaction modalities in single-cell perturbation experiments.

#### Note S9: Model evaluation for predicting genetic interactions

Following the evaluation framework established by GEARS, the PRJNA551220 scCRISPR-seq dataset generated by Norman et al. was utilized to assess scPert's performance in predicting genetic interactions. After quality control and preprocessing, 125 dual genetic perturbations were retained for analysis. To evaluate the model's ability to accurately predict different GI subtypes including synergy, suppression, neomorphism, redundancy, and epistasis (Note S6). Specifically, for each of the 125 two-gene combinatorial perturbations experimentally assayed in the Norman et al. study, we trained scPert from scratch while holding out that specific interaction as the test set. This rigorous evaluation strategy ensures that the model must generalize to completely unseen genetic interactions during testing.

Once each model completed training, genetic interaction scores are calculate for the held-out perturbation in the test dataset. These GI scores quantify different aspects of combinatorial gene effects: magnitude scores measure the overall strength of the interaction, model fitting scores assess deviation from additive effects, equality of contribution scores evaluate whether both genes contribute equally to the phenotype, and similarity scores compare the combinatorial effect to individual gene effects. The single GI score threshold system are utilized for identifying the GI subtype proposed by Roohani et al., which categorizes interactions based on predefined cutoffs for each score metric (Note S8).

Model performance in predicting each GI subtype was evaluated using two key metrics. The Precision@10 metric quantifies the fraction of true positives among the top 10 predicted interactions for each subtype, measuring the model's ability to prioritize the most likely genetic interactions for experimental validation. This metric is particularly valuable for guiding targeted experimental follow-up, as it indicates how many of the highest-confidence predictions would be validated if subjected to experimental testing (Note S7). The Accuracy metric, evaluated using our decision tree-based threshold system, measures the overall proportion of correctly classified interactions across all five interaction types, providing a comprehensive assessment of the classification performance of methods across the full spectrum of genetic interaction phenotypes.

#### Note S10: Performance evaluation on p53 pathway perturbations

The systematic evaluation employed three complementary metrics: Pearson correlation measuring expression relationship accuracy, direction accuracy assessing up/down-regulation prediction correctness, and MAE quantifying magnitude prediction error. scPert demonstrated superior overall performance with the highest Pearson correlation (0.927) and lowest MAE (0.153), indicating excellent capture of gene expression relationships and quantitative accuracy following p53 pathway perturbations. While scGPT achieved the highest direction accuracy (0.820), scPert's combined performance across correlation and magnitude prediction, coupled with competitive direction accuracy (0.790), establishes it as the most effective method for p53 pathway analysis. GEARS showed intermediate performance across all metrics, suggesting balanced but suboptimal capabilities for this clinically important tumor suppressor pathway (detailed results in **Table S8**).

#### Note S11: Mixscape-based quality control and perturbation validation

High-throughput single-cell perturbation experiments require rigorous quality control measures to ensure accurate identification of successfully perturbed cells and removal of cells that escaped the intended perturbation (i.e., "escape cells" that retain wild-type expression of the target gene despite guide RNA delivery). We employed Mixscape[10], a computational method specifically designed for single-cell CRISPR screen analysis, and followed its official documentation for all analytical procedures to systematically evaluate perturbation efficiency and filter out escape cells from our datasets. Mixscape leverages Bayesian probabilistic models to classify cells into "perturbed" or "unperturbed/escape" categories by integrating three key layers of information: (1) guide RNA (gRNA) expression levels, (2) target gene knockdown efficiency, and (3) global transcriptional changes consistent with successful perturbation.

The application of Mixscape to our perturbation datasets revealed substantial heterogeneity in perturbation success rates across different experimental conditions and target genes. In the immune checkpoint regulation dataset containing THP-1 cells (human monocytic leukemia line), we initially began with 99 different perturbation conditions targeting various regulatory genes involved in PD-L1 expression control. Following Mixscape-based quality filtering, we identified only 11 perturbation conditions that met stringent criteria for successful perturbation, representing an approximately 89% reduction in the dataset size. This dramatic filtering reflects the technical challenges inherent in CRISPR-based perturbation experiments, particularly in primary immune cells where perturbation efficiency can be variable due to cellular heterogeneity, guide RNA delivery issues, and intrinsic resistance to genetic modification.

#### Note S12: Prediction quality score calculation methodology

To comprehensively evaluate scPert's prediction accuracy for cancer target gene analysis, we developed a composite quality scoring system that assesses both directional consistency and numerical precision between predicted and experimental $\log_2$ fold changes ($Log_2FC$). The prediction quality score (QS) is calculated as a weighted combination of directional accuracy and numerical proximity using the formula: $QS = 0.6 \times \text{Direction Score} + 0.4 \times \text{Proximity Score}$.

The direction score component evaluates whether the predicted perturbation effect matches the experimental direction. For predicted $Log_2FC$ (P) and true $Log_2FC$ (T), the direction score is assigned as follows: when both $|P|$ and $|T|$ exceed the threshold of 0.1, the direction score equals 1.0 if $\text{sign}(P) = \text{sign}(T)$, and 0.0 if the signs differ. When only one value exceeds the threshold (indicating one directional and one neutral response), the direction score is set to 0.5. When both values fall below the threshold (both neutral), the direction score is 1.0. This threshold of 0.1 $Log_2FC$ was empirically determined to balance sensitivity and specificity for identifying meaningful expression changes.

The proximity score measures numerical accuracy using an exponential decay function: $\text{Proximity Score} = \exp(-|P - T| \times 2)$. This formulation ensures that perfect predictions ($P = T$) receive a score of 1.0, while predictions with larger absolute differences receive exponentially lower scores. The decay factor of 2 provides appropriate sensitivity to prediction errors, creating a smooth transition between high and low accuracy predictions.

Quality scores are subsequently categorized into five levels: Excellent ($QS \geq 0.8$) indicating high directional and numerical accuracy, Good ($0.6 \leq QS < 0.8$) representing good overall performance, Fair ($0.4 \leq QS < 0.6$) denoting moderate accuracy, Poor ($0.2 \leq QS < 0.4$) indicating low accuracy, and Very Poor ($QS < 0.2$) representing very low accuracy. The sixty-forty weighting scheme prioritizing directional accuracy over numerical precision is based on the biological principle that regulatory direction determines functional consequences in gene regulatory networks. Whether a gene is activated or repressed by perturbation fundamentally affects its downstream impact on cellular phenotypes, making directional consistency more critical than precise magnitude quantification for most biological applications.

#### Note S13: CompGCN integration and ablation analysis

Knowledge graph embedding (KGE) methods are fundamental for encoding structured biological relationships, with traditional approaches including relation-agnostic standard graph neural networks and translation-based models like TransE. Standard GCNs treat all edges in biological networks uniformly, failing to capture the distinct semantics of different relationship types, while TransE constrains interactions to simple vector translations that limit modeling of complex compositional patterns. These limitations highlight the need for advanced KGE architectures tailored to heterogeneous biological networks.

The integration of Compositional Graph Convolutional Networks (CompGCN)[11] within the scPert framework demonstrates the critical importance of advanced graph neural network architectures for modeling complex gene regulatory relationships in combinatorial perturbation scenarios. CompGCN extends traditional graph convolutional networks by jointly embedding both nodes and edge types through compositional operations, enabling more sophisticated representation of heterogeneous biological networks where genes interact through diverse regulatory mechanisms.

Ablation analysis across different generalization scenarios reveals the substantial contribution of CompGCN to scPert's superior performance, particularly in challenging combinatorial perturbation prediction tasks. In the most difficult scenario (seen0, corresponding to 0/2 unseen where neither gene in the combination had been individually perturbed during training), scPert with CompGCN achieved an MSE of 0.120, representing a significant 62% improvement over the basic graph variant (0.314) and demonstrating competitive performance against established methods like GEARS (0.132) and scGPT (0.143). The improvement becomes even more pronounced in intermediate difficulty scenarios, with CompGCN achieving 0.167 in seen1 (1/2 unseen) compared to 0.259 for the graph-only version, and 0.078 in seen2 (2/2 seen) versus 0.212 for the basic graph implementation.

The superior performance of CompGCN stems from its ability to learn compositional relationships between different edge types in biological networks, capturing the complex interplay between transcriptional regulation, protein interactions, and metabolic dependencies that govern cellular responses to genetic perturbations. Unlike standard graph neural networks that treat all connections uniformly, CompGCN's compositional framework enables nuanced modeling of how different relationship types contribute to perturbation effects, making it particularly effective for extrapolating to novel gene combinations where regulatory principles must be inferred from learned compositional rules rather than direct experimental observations.

To further validate the role of structured biological knowledge, we conducted ablation experiments removing the Hetionet knowledge graph component (while retaining foundation model embeddings and hierarchical fusion). Centroid accuracy decreased substantially across all five datasets: from 0.72 to 0.56 (Adamson), 0.66 to 0.59 (Norman), 0.60 to 0.50 (Dixit, reaching random baseline), 0.75 to 0.52 (Replogle K562), and 0.68 to 0.50 (Replogle RPE1, random baseline). These results confirm Hetionet's essential role in providing biological constraints to distinguish perturbation-specific signals from noise, with detailed data in **Table S10**.

#### Note S14: Loss function of scPert

The scPert model employs a multi-component loss function designed to capture multiple aspects of gene expression perturbation dynamics. The total loss function combines five complementary objectives computed separately for each perturbation before averaging to ensure balanced gradient contributions. The focal MSE loss automatically emphasizes genes showing larger perturbation-induced changes through adaptive weighting, providing moderate focus on differentially expressed genes. The weighted MSE loss explicitly incorporates biological prior knowledge by assigning higher importance to specified gene sets such as transcription factors and key signaling molecules, where class weights boost important genes by a factor of 5 while maintaining normalized weights to prevent numerical instability. The direction loss explicitly penalizes predictions with incorrect regulatory directions, ensuring the model captures not just expression magnitudes but also up-regulation and down-regulation patterns critical for understanding gene regulatory relationships. The cosine similarity loss provides global constraints on expression profiles, preserving co-expression patterns and regulatory modules that are invariant to magnitude scaling. Optional L1 regularization promotes parameter sparsity, though we found that careful architecture design combined with other loss components typically provides sufficient regularization.

$Total\ Loss\ Function$:

$$\mathcal{L}total = \mathcal{L}focal + \mathcal{L}weighted + \lambda dir\mathcal{L}direction + \lambda cos\mathcal{L}cosine + \lambda l1\mathcal{L}reg$$

The loss function incorporates several design features critical for robust perturbation prediction. Perturbation-specific gene filtering allows computing losses only on genes relevant to each perturbation, improving computational efficiency and enabling incorporation of pathway-specific knowledge. Our hyperparameter settings were selected through systematic grid search on validation data and provide robust performance across diverse datasets.

This multi-objective loss formulation enables scPert to learn robust representations that capture both local gene-specific dynamics and global expression patterns while incorporating biological constraints through weighted MSE and directional supervision. The focal mechanism adaptively emphasizes highly perturbed genes without requiring manual specification of importance, while the weighted component allows explicit incorporation of domain knowledge about biologically critical regulators. The direction loss ensures preservation of regulatory relationships essential for understanding causal gene networks, and the cosine similarity loss maintains overall expression profile structure across predictions. By balancing these complementary objectives through carefully tuned hyperparameters, the loss function provides comprehensive supervision that enables scPert to generalize effectively to unseen perturbations, handle combinatorial genetic interactions, and achieve state-of-the-art performance across diverse perturbation prediction benchmarks. The perturbation-specific filtering mechanism further enhances efficiency and enables pathway-focused predictions when relevant, making the loss function adaptable to various experimental designs and biological questions while maintaining robust training dynamics through per-perturbation computation and normalized weighting schemes.

#### Note S15: Hyperparameters of scPert

The scPert model was developed using the PyTorch framework and all experiments of this study were performed on a machine with one NVIDIA A100 GPU and 80GB memory. The model architecture integrates multiple sources of biological knowledge through a multi-modal embedding strategy combined with transformer-based processing. Specifically, scPert incorporates three types of embeddings: knowledge graph embeddings (KGE) generated from biological networks using CompGCN (128-dimensional), contextualized gene embeddings from the pre-trained scGPT foundation model (512-dimensional), and learnable gene-specific representations. For perturbation genes, we concatenate their KGE and scGPT embeddings to form 640-dimensional combined representations, which are then projected to the model's hidden dimension of 64 through dense layers with LayerNorm, ReLU activation, and dropout rate of 0.1.

The core architecture consists of an embedding fusion module, a multi-layer transformer encoder, and a gene-specific prediction head. The embedding fusion module processes gene and perturbation information through separate pathways. Gene embeddings are first projected to 64 dimensions and enhanced with a gene interaction layer that computes pairwise attention scores between all genes to capture gene-gene relationships. Perturbation embeddings (640-dimensional) are processed through a two-layer MLP with intermediate dimension of 128, reducing to 64 dimensions. For combination perturbations involving multiple genes, individual perturbation embeddings are fused using mean pooling after MLP transformation. The transformer encoder consists of 8 stacked layers with 8-head self-attention. To handle large sequence lengths (thousands of genes), we implemented a memory-efficient chunked attention mechanism with chunk size of 256. Each layer uses GELU activation in feed-forward networks with an expansion factor of 4, layer normalization in pre-norm configuration, and residual connections with 0.5 scaling factors. The final prediction head comprises three fully-connected layers (64 → 256 → 128 → 64 dimensions) with LayerNorm, ReLU, and dropout, followed by a linear output layer projecting to predicted gene expression changes.

Model parameters were optimized using the AdamW optimizer with learning rate of 0.002 and weight decay of $1\times10^{-5}$. We employed OneCycleLR scheduler with maximum learning rate of 0.01, implementing cyclical learning rate policy with cosine annealing. Gradient clipping with maximum norm of 1.0 was applied to prevent gradient explosion. The model was trained for 20 epochs with batch size of 32.

### Supplementary Figures

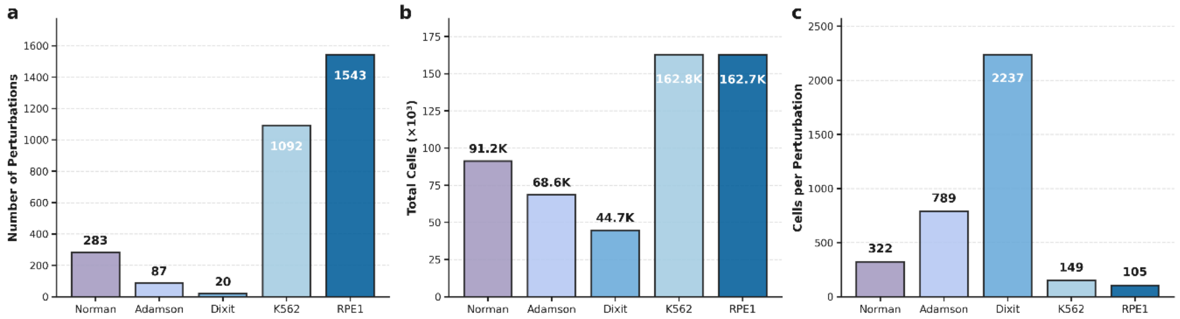


**Figure S1. Statistical overview of perturbation datasets used for model evaluation.** (*a*) Number of perturbations ranging from 20 to 1,543 across five datasets. (*b*) Total cell counts from 44.7K to 162.8K cells per dataset. (*c*) Cells per perturbation ratio varying from 105 to 2,237, reflecting distinct experimental design strategies.

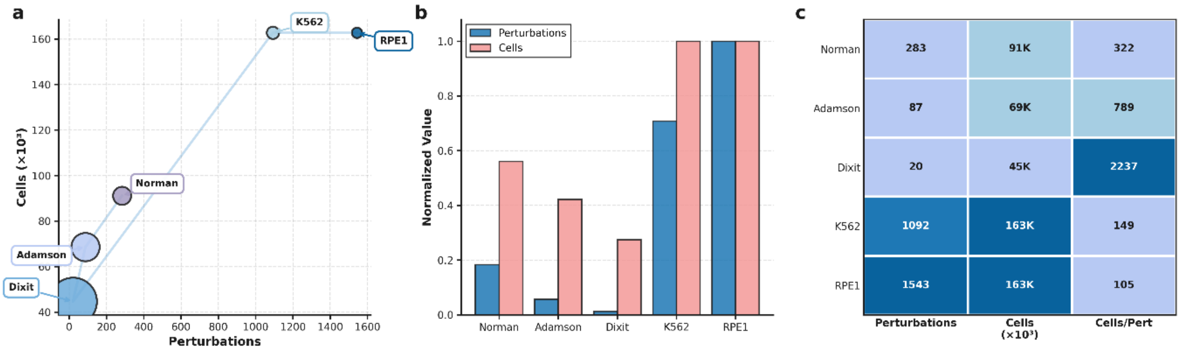


**Figure S2. Detailed statistical characterization of benchmark datasets** (***a***) Scatter plot showing the relationship between perturbations and total cells. Bubble size represents cells per perturbation. (***b***) Normalized comparison of perturbations (blue) and cells (pink) across datasets. (***c***) Heatmap displaying normalized intensities for perturbation count, total cells, and cellular sampling depth.

***a**. Prediction of Top 20 DE Genes in Single Perturbation*

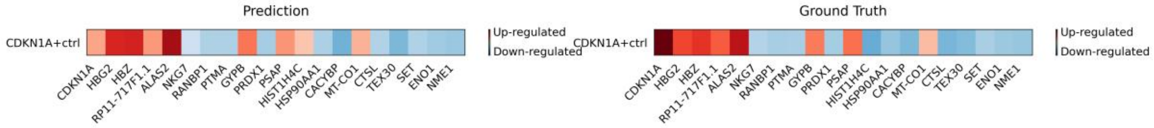


***b**. Prediction of Top 20 DE Genes in Combination Perturbations*

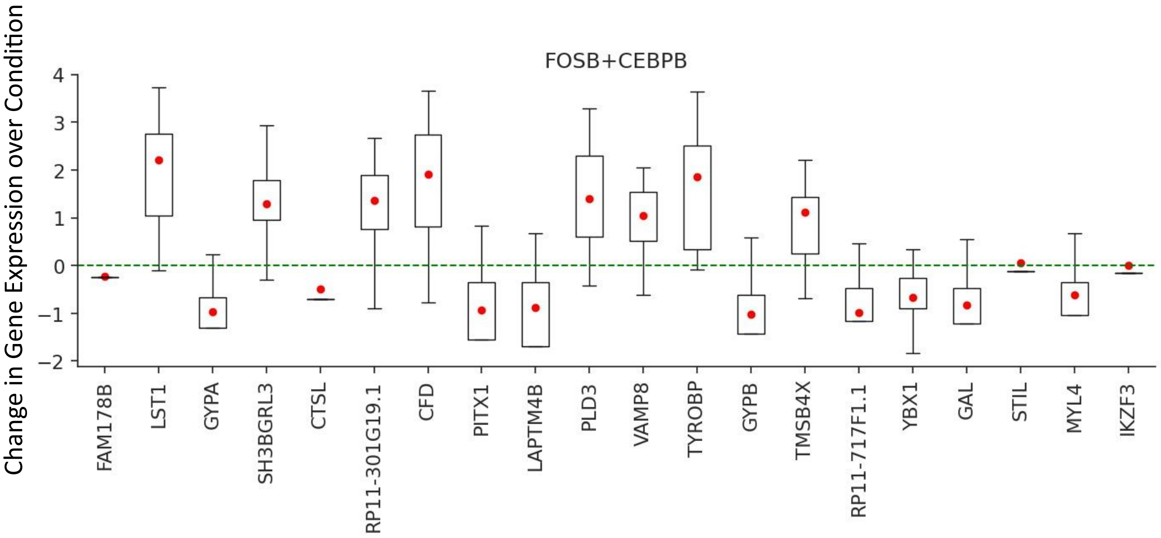


**Figure S3. Prediction of scPert for Top 20 DE genes in different perturbations.** (***a***) Heatmap comparison of predicted versus ground truth gene expression changes for the top 20 differentially expressed genes following CDKN1A+ctrl single perturbation. Red indicates upregulation and blue indicates downregulation. The model accurately captures both the identity and directionality of most differentially expressed genes. (***b***) Box plot showing predicted gene expression changes for the top 20 differentially expressed genes in FOSB+CEBPB combination perturbation. Each box represents the distribution of predicted expression changes across multiple cells, with red dots indicating the median values. The horizontal dashed line at zero represents no expression change. The results demonstrate scPert's capability to predict complex non-additive gene expression patterns that emerge from combinatorial perturbations, capturing both synergistic upregulation and suppressive effects across different functional gene categories.

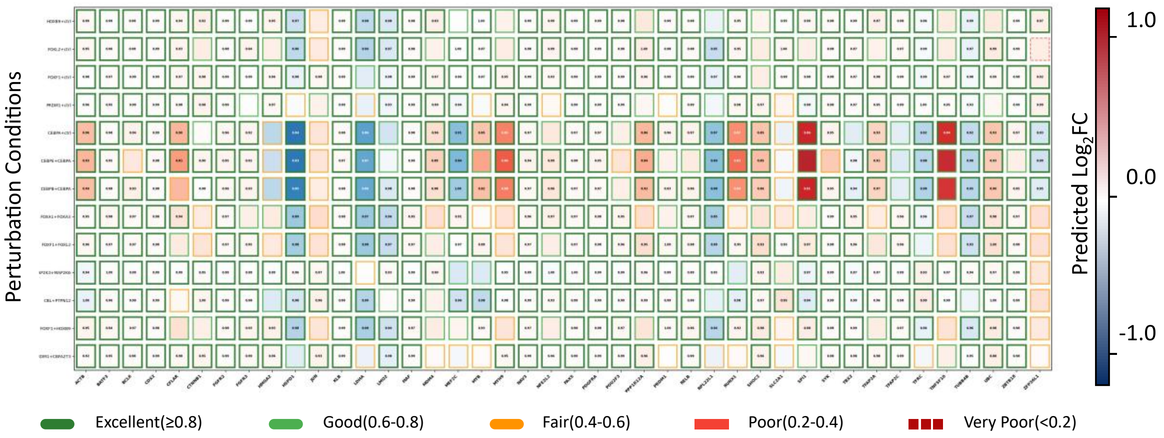


**Figure S4. Comprehensive evaluation of scPert prediction quality across perturbation conditions and target genes.** The heatmap displays prediction quality scores for different perturbation conditions (y-axis) against target genes (x-axis), with colors representing predicted $Log_2FC$ values. Quality scores are categorized as Excellent (≥0.8, dark green), Good (0.6-0.8, light green), Fair (0.4-0.6, orange), Poor (0.2-0.4, red), and Very Poor (<0.2, dark red) based on prediction accuracy. The results demonstrate that scPert achieves excellent or good prediction quality for the majority of perturbation-gene combinations, with some challenging cases showing lower prediction accuracy. This systematic evaluation reveals the model's robust performance across diverse perturbation conditions while identifying specific gene-perturbation combinations that may require additional optimization or represent inherently difficult prediction scenarios.

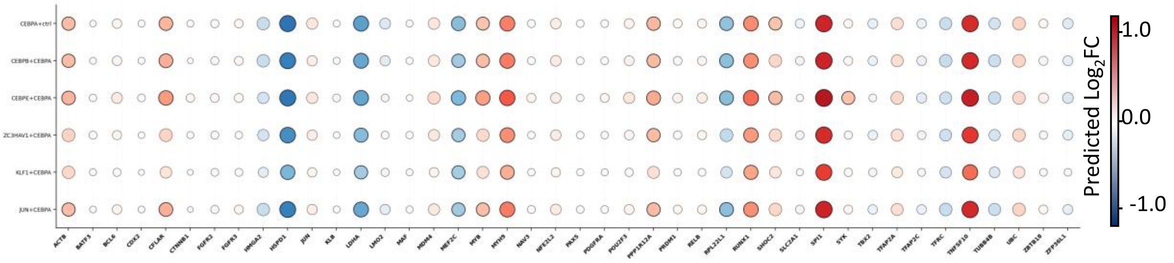


**Figure S5**. **CEBPA-dominant perturbation effects on gene expression.** The vertical axis represents different perturbation conditions, and the horizontal axis shows target genes. Bubble color and size reflect the magnitude of predicted gene expression changes ($Log_2FC$). The results demonstrate that combination perturbations involving CEBPA (CEBPA+CEBPE, CEBPA+CEBPB and others) exhibit remarkably similar expression profiles to CEBPA single perturbation across the target gene set, confirming CEBPA's dominant regulatory effects in these combinatorial contexts, where its transcriptional program overrides the effects of co-perturbed transcription factors.

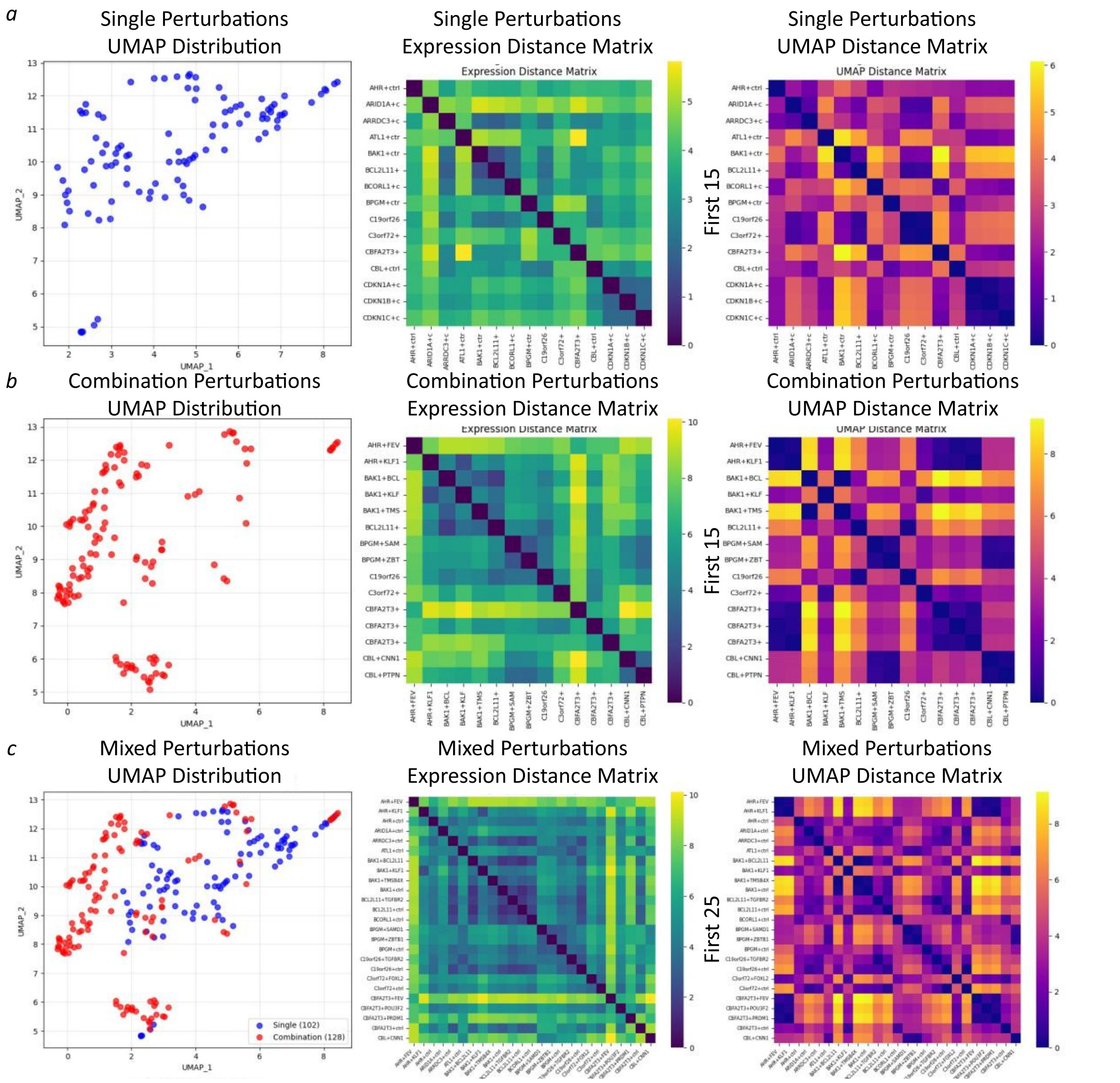


**Figure S6. UMAP representation and distance correlation analysis of single versus combination perturbations.** (***a***) Single Perturbations. Spatial distribution and distance relationships of single-gene perturbation conditions in UMAP space. Left panel shows the distribution of single-gene perturbations in UMAP coordinates (blue dots); middle panel displays the expression distance matrix heatmap based on Euclidean distances calculated from gene expression profiles; right panel shows the corresponding UMAP distance matrix heatmap. Colors range from dark blue/purple (short distances) to yellow (long distances). (***b***) Combination Perturbations Spatial distribution and distance relationships of combination perturbation conditions (involving two or more genes) in UMAP space. Left panel shows the distribution of combination perturbations in UMAP coordinates (red dots); middle and right panels show expression distance and UMAP distance matrix heatmaps, respectively. Combination perturbations exhibit distinct clustering patterns compared to single perturbations. (***c***) Cross-Type Comparison. Comparative analysis of single-gene perturbations (blue) versus combination perturbations (red) in the same UMAP space and their cross-type distance relationships. Left panel shows the overlay of both perturbation types in UMAP space; middle panel displays the expression distance matrix between single and combination perturbations; right panel shows the corresponding UMAP distance matrix. This analysis reveals the spatial separation and interaction patterns between different perturbation types.

### Supplementary Tables

**Table S1.** Statistical overview of perturbation datasets used for model evaluation

| Statistic | Norman | Adamson | Dixit | Replogle K562 | Replogle RPE1 |
|---|---|---|---|---|---|
| Pert | 283 | 87 | 20 | 1092 | 1543 |
| Cell | 91205 | 68603 | 44735 | 162751 | 162733 |

**Table S2**. Mean squared error performance of single gene perturbation

| Method | Norman | Adamson | Dixit | Replogle K562 | Replogle RPE1 |
|---|---|---|---|---|---|
| Gears | 0.216 | 0.242 | 0.088 | 0.147 | 0.152 |
| scGPT | 0.239 | 0.169 | 0.106 | 0.154 | 0.131 |
| scGenePT (ncbi+uniprot) | 0.215 | 0.144 | — | — | — |
| scGenePT (go_all) | 0.214 | 0.145 | — | — | — |
| Biolord | 0.215 | 0.212 | 0.236 | 0.148 | 0.145 |
| Scouter | 0.321 | 0.136 | 0.013 | 0.122 | 0.150 |
| scPert | 0.190 | 0.151 | 0.091 | 0.123 | 0.122 |

*Note: It should be noted that scGenePT comparisons were limited to the Norman and Adamson datasets, as only checkpoints for these two benchmarks are currently publicly available from the original authors. We evaluated both the ncbi+uniprot and go_all model configurations to ensure comprehensive comparison with scGenePT's top-performing variants. Complete benchmarking across all five datasets was not feasible for scGenePT at the time of this study.*

**Table S3**. Pearson correlation performance of single gene perturbation

| Method | Norman | Adamson | Dixit | Replogle K562 | Replogle RPE1 |
|---|---|---|---|---|---|
| Gears | 0.990 | 0.958 | 0.921 | 0.904 | 0.658 |
| scGPT | 0.991 | 0.957 | 0.932 | 0.905 | 0.696 |
| scGenePT (ncbi+uniprot) | 0.987 | 0.978 | — | — | — |
| scGenePT (go_all) | 0.987 | 0.976 | — | — | — |
| Biolord | 0.944 | 0.965 | 0.896 | 0.989 | 0.636 |
| Scouter | 0.897 | 0.969 | 0.996 | 0.971 | 0.668 |
| scPert | 0.988 | 0.962 | 0.912 | 0.897 | 0.683 |

*Note: It should be noted that scGenePT comparisons were limited to the Norman and Adamson datasets, as only checkpoints for these two benchmarks are currently publicly available from the original authors. We evaluated both the ncbi+uniprot and go_all model configurations to ensure comprehensive comparison with scGenePT's top-performing variants. Complete benchmarking across all five datasets was not feasible for scGenePT at the time of this study.*

**Table S4**. Centroid accuracy performance of single gene perturbation

| Method | Norman | Adamson | Dixit | Replogle K562 | Replogle RPE1 |
|---|---|---|---|---|---|
| Gears | 0.602 | 0.480 | 0.650 | 0.499 | 0.529 |
| scGPT | 0.610 | 0.510 | 0.570 | 0.550 | 0.520 |
| scGenePT (ncbi+uniprot) | 0.703 | 0.576 | — | — | — |
| scGenePT (go_all) | 0.703 | 0.608 | — | — | — |
| Biolord | 0.671 | 0.660 | 0.500 | 0.652 | 0.640 |
| Scouter | 0.400 | 0.320 | 0.210 | 0.727 | 0.640 |
| scPert | 0.663 | 0.724 | 0.600 | 0.754 | 0.684 |
| Non control mean | 0.533 | 0.500 | 0.500 | 0.500 | 0.500 |

*Note: It should be noted that scGenePT comparisons were limited to the Norman and Adamson datasets, as only checkpoints for these two benchmarks are currently publicly available from the original authors. We evaluated both the ncbi+uniprot and go_all model configurations to ensure comprehensive comparison with scGenePT's top-performing variants. Complete benchmarking across all five datasets was not feasible for scGenePT at the time of this study.*

**Table S5**. Mean squared error performance of combinatorial perturbation

| Method | Seen 0 | Seen 1 | Seen 2 |
|---|---|---|---|
| Gears | 0.132 | 0.177 | 0.180 |
| scGPT | 0.143 | 0.170 | 0.133 |
| scPert | 0.121 | 0.167 | 0.078 |

**Table S6**. Pearson correlation performance of combinatorial perturbation

| Method | Seen 0 | Seen 1 | Seen 2 |
|---|---|---|---|
| Gears | 0.655 | 0.591 | 0.625 |
| scGPT | 0.647 | 0.601 | 0.660 |
| scPert | 0.693 | 0.652 | 0.736 |

**Table S7.** Precision@10 scores for genetic interaction prediction

| Method | Synergy | Suppressive | Neomorphism | Redundant | Epistasis |
|---|---|---|---|---|---|
| Gears | 0.60 | 0.60 | 0.50 | 0.9 | 0.90 |
| scGPT | 0.80 | 0.80 | 0.60 | 1.0 | 1.0 |
| scPert | 0.90 | 1.0 | 0.60 | 0.9 | 1.0 |

**Table S8.** Performance evaluation on p53-associate gene

| Method | Pearson correlation | Direction Accuracy | MAE |
|---|---|---|---|
| Gears | 0.899 | 0.690 | 0.169 |
| scGPT | 0.907 | 0.820 | 0.191 |
| scPert | 0.927 | 0.790 | 0.153 |

**Table S9**. Mean squared error performance of ablation experiment

| Scenario | Gears | scGPT | scPert_Graph | scPert_TransE | scPert_CompGCN |
|---|---|---|---|---|---|
| Seen 0 | 0.132 | 0.143 | 0.314 | 0.262 | 0.120 |
| Seen 1 | 0.177 | 0.170 | 0.259 | 0.274 | 0.167 |
| Seen 2 | 0.180 | 0.133 | 0.212 | 0.155 | 0.078 |

**Table S10**. Ablation experiments on centroid accuracy

| Method | Norman | Adamson | Dixit | Replogle K562 | Replogle RPE1 |
|---|---|---|---|---|---|
| scPert_No_KG | 0.591 | 0.557 | 0.500 | 0.522 | 0.504 |
| Non control mean | 0.533 | 0.500 | 0.500 | 0.500 | 0.500 |

### Reference


1. Norman, T.M. et al. Exploring genetic interaction manifolds constructed from rich single-cell phenotypes. *Science* **365**, 786-793 (2019).
2. Adamson, B. et al. A Multiplexed Single-Cell CRISPR Screening Platform Enables Systematic Dissection of the Unfolded Protein Response. *Cell* **167**, 1867-1882.e1821 (2016).
3. Replogle, J.M. et al. Mapping information-rich genotype-phenotype landscapes with genome-scale Perturb-seq. *Cell* **185**, 2559-2575.e2528 (2022).
4. Dixit, A. et al. Perturb-Seq: Dissecting Molecular Circuits with Scalable Single-Cell RNA Profiling of Pooled Genetic Screens. *Cell* **167**, 1853-1866.e1817 (2016).
5. Roohani, Y., Huang, K. & Leskovec, J. Predicting transcriptional outcomes of novel multigene perturbations with GEARS. *Nature biotechnology* **42**, 927-935 (2024).
6. Cui, H. et al. scGPT: toward building a foundation model for single-cell multi-omics using generative AI. *Nat Methods* **21**, 1470-1480 (2024).
7. Piran, Z., Cohen, N., Hoshen, Y. & Nitzan, M. Disentanglement of single-cell data with biolord. *Nat Biotechnol* **42**, 1678-1683 (2024).
8. Istrate, A.-M., Li, D. & Karaletsos, T. scGenePT: Is language all you need for modeling single-cell perturbations? *bioRxiv* (2024). https://doi.org/10.1101/2024.10.23.619972
9. Zhu, O. & Li, J. Scouter predicts transcriptional responses to genetic perturbations with large language model embeddings. *Nat Comput Sci* (2025). https://doi.org/10.1038/s43588-025-00912-8
10. Papalexi, E. et al. Characterizing the molecular regulation of inhibitory immune checkpoints with multimodal single-cell screens. *Nature genetics* **53**, 322-331 (2021).
11. Vashishth, S., Sanyal, S., Nitin, V. & Talukdar, P. Composition-based Multi-Relational Graph Convolutional Networks. in International Conference on Learning Representations (2020).
